# Convolvulus pluricaulis mediates its pharmacological effects via sod1, rdl, glut1, GABA-B-R1 and CG6293 orthologs in Drosophila melanogaster

**DOI:** 10.1101/2025.07.06.663350

**Authors:** Shreyasi Mitra, Amit Kumar, Manish Pandey, Meenakshi Sharma, Shivani Pundir, Mansi Jangir, Geetanjali Chawla

## Abstract

*Convolvulus pluricaulis,* commonly known as Shankhpushpi, has been extensively used in the management of disorders of the nervous system, including depression and anxiety. The plant extract has also been demonstrated to function as an antioxidant. However, the molecular effectors underlying the beneficial effects of Shankpushpi are yet to be identified. In this study, we have utilized the fruit fly *Drosophila melanogaster* to identify the metabolic and molecular targets that mediate the beneficial effects of dietary Shankpushpi intake. Metabolomic analysis revealed changes in Ascorbic acid, glucose, and Adenosine monophosphate in the head tissue of fruit flies that were fed Shankpushpi for 20 days. Subsequent gene expression analysis revealed significant changes in expression of *glut 1* (Glucose transporter 1), *CG6293* (Ascorbate transporter), *rdl* (Resistant to dieldrin*), GABA-B-R1* (GABA-B receptor subtype 1) and *sod 1* (Superoxide dismutase 1) in the head tissue of adult flies that were exposed to different doses of Shankpushpi. Consistent with the expression data, knockdown of *sod 1*, *glut1*, *GABA-B-R1*, and *CG6293* but not *sod 2* abolished the Shankpushpi-mediated resistance to paraquat-induced oxidative stress. To uncover downstream effectors that are responsible for the antidepressant and anxiolytic effects of Shankpushpi, we examined the effects of dietary intake of Shankpushpi in a stress-induced model of depression. Administration of Shankpushpi during development alleviated depression-like behavior in *Drosophila*. Wild-type adult flies that were fed Shankpushpi accumulated high levels of Ascorbate in the head tissue, and knockdown of Ascorbate transporter abolished the antidepressant activity of Shankpushpi in the stress-induced model. Thus, indicating that Shankpushpi-induced antidepressant effects are associated with increased Ascorbate transport. Taken together, our analysis using *Drosophila melanogaster* as a model has uncovered five conserved downstream effectors responsible for the antioxidant activity and one conserved effector responsible for the antidepressant activity of *Convolvulus pluricaulis*.

## 1. INTRODUCTION

Depression and anxiety are recurrent mental health disorders that severely affect the patient’s quality of life due to frequent relapses and remissions[1–3]. The symptoms of depression include behavioral despair, anhedonia, helplessness, fatigue, and sleep disturbances. Some of the hypotheses proposed for the pathogenesis of depression include abnormal expression of neurotransmitters and their receptors, deregulation of the immune system and inflammation, and oxidative stress dysfunction of the hypothalamic-pituitary axis, reduction in the levels of neurotrophic factors, dysregulation of gut microbiota, mitochondrial dysfunction and a reduction in neuroplasticity impacting neuronal connections [4–11].

Pharmacological medications that are used for depression and anxiety include selective serotonin reuptake inhibitors (SSRIs), Serotonin-Noradrenaline reuptake inhibitors (SNRIs), Atypical antidepressants, Serotonin-Dopamine Activity modulators (SDAMs), Tricyclic antidepressants (TCAs) and monoamine oxidase inhibitors [12]. These medications are not effective in the long term for all individuals and can cause side effects. Moreover, most of these drugs provide temporary relief and are not curative. Natural plant-based supplements are increasingly being considered for promoting health and treat mental disorders with minimal side effects[13, 14]. These are complex phytochemical mixtures that contain one or more bioactive compounds, providing a single plant extract with multiple beneficial properties[15, 16].

*Convovulus pluricaulis* is a perennial plant that has been traditionally used for its neuropharmacological benefits, including but not limited to antidepressant, antioxidant, memory-enhancing, and anxiolytic effects[17–24]. Some of the broad classes of phytoconstituents that have been identified in *C. pluricaulis* (Shankhpushpi) include Vitamin E, ascorbic acid, phthalic acid, squalene, silane, decanoic acid, linoleic acid, tropane alkaloids, kempferol, beta-sitosterol, pentanoic acid, and cinnamic acid [25–27]. Despite the availability of phytochemical information regarding the constituents of this medicinal extract, the molecular targets of these constituents remain largely unknown.

In this study, we utilized *Drosophila melanogaster* as a model to uncover the molecular effectors underlying the anti-depressant and antioxidant effects of *Convovulus pluricaulis* (Shankhpushpi). We first set up a variable stress-induced *Drosophila* model to assess the impact of *C. pluricaulis* intake on depression like behavior. Our studies revealed that *C. pluricaulis* )(Shankhpushpi) alleviates anhedonia in a dose-dependent manner in both male and female flies. To identify tissue-specific changes in the metabolome, we performed GC-MS analysis on the head and body tissues of wild-type flies fed a control or Shankhpushi-supplemented diet. This analysis revealed a significant increase in L-Ascorbate and D-Glucose in the head tissue of flies that were fed Shankhpushpi. Consistent with the metabolomic data, we observed a significant increase in expression of *L-Ascorbate transporter* (*CG6293*), *Glucose transporter 1* (*Glut1*), and the oxidative stress defence gene *Superoxide dismutase 1* (*Sod1*). Increase in Oxidative stress and a decreased levels of antioxidative enzymes have been associated with neuropsychiatric diseases[28, 29]. Due to its higher metabolic rate, higher oxidizable lipid content, and less robust antioxidant defense, brain tissue is more vulnerable to oxidative stress[30].

Exhaustion of antioxidative defenses leads to the excessive generation of reactive oxygen species (ROS) that trigger proinflammatory signaling and damage to cellular macromolecules. Patients suffering from depression have significantly lower plasma levels of antioxidants such as Vitamin E, zinc, and coenzyme Q_10_ (CoQ_10_), along with a lower antioxidant capacity[31]. Non-enzymatic antioxidants (reduced glutathione or GSH) and antioxidant enzymes such as Superoxide dismutase together contribute the role of oxidative stress in depression-like states. To examine whether *CG6293* or *Sod1* was contributing to the antidepressant activity of *C. pluricaulis*, we tested the ability of the *C. pluricaulis* to alleviate stress-induced anhedonia after ubiquitously knocking down *CG6293* and *Sod1*. We tested whether reducing the levels of these genes compromised the anti-depressant activity of *C. pluricaulis* (Shankhpushpi). Knockdown of *CG6293* but not *Sod1* in adult flies eliminated the resilience provided by Shankhpushpi in the stress-induced depression model.

Next, we tested whether reducing the levels of the Shankhpushpi-modulated genes resulted in alleviation of its antioxidant property. The ability of Shankhpushpi to enhance survivability of fruit flies exposed to paraquat-induced oxidative stress was measured after reducing the dosage of *Sod1*, *GABA-B-R1*, *Rdl*, *Sod2*, *Glut1,* and *CG6293*. Reducing one copy of *GABA-B-R1*, *Rdl*, *Glut1*, *CG6293,* and *Sod1* or ubiquitous knockdown of *sod1* resulted in no significant increase in lifespan in the presence of Shankhpushpi. In contrast, knockdown of *Sod2* did not alter the ability of Shankhpushpi to enhance survivability of flies that were exposed to Paraquat. Thus, indicating that Shankhpushpi mediates it antioxidant property via sod1, CG6293, GABA-B-R1, Glut1 and Rdl. Overall, our analyses have identified conserved downstream effectors that are responsible for the antidepressant and antioxidant properties of *C. pluricaulis*.

## 2. METHODS

### 2.1. *Drosophila* strains and maintenance

*Drosophila melanogaster* stocks of the wildtype strain *Canton* S and *w^1118^* have been in the laboratory since 2017 and maintained in a sugar-yeast cornmeal-based diet (please see section 2.2 for recipe). *Canton S* flies were used for experiments depicted in Figure 1, Figure 2, Figure 3, Figure 5, and Supplementary Figure 1. The *UAS-Sod2 ^RNAi^* line (RRID: BDSC_36871), *UAS-Sod1^RNAi^* (RRID: BDSC_24491), *UAS-CG6293^RNAi^* (VDRC108619), *Rdl*[1] (RRID: BDSC_1687), GABA-B-R1 mimic line (RRID: BDSC_36226), *Glut1* mimic line (RRID: BDSC_37890), CG6293 mimic line (RRID: BDSC_59511), *sod1* mimic line (RRID: BDSC_44929), and *Da-GS* Gal4 (gift from David Walker’s laboratory) driver were used for the experiments in Figure 6. The Da-GS lines and UAS lines used for lifespan were backcrossed three times into a homogenous control background *w^1118^* by setting up crosses with a single male from each line with three female *w^1118^* virgin flies. The process was repeated with F2 males crossed to *w^1118^* virgin flies, F3 males crossed to *w^1118^* virgin flies, and the F4 male progeny were crossed with a compound chromosome 2:3 lab balancer stock. The resulting balanced stocks were used for setting up crosses for lifespan analysis. All flies used in the study were maintained in standard cornmeal/agar medium at 25°C with a 12h light: 12h dark cycle in 60% humidity. For steroid-mediated knockdown of *Sod1* and *Sod2* using the Gene-Switch driver, flies were fed a diet containing 200μM RU-486 (Mifepristone, Cayman Chemicals, Ann Arbor, MI). Comparisons of lifespan were made in parallel with the same strain on different diets (with and without Shankhpushpi). The lack of any effect of RU-486 on *Da-GS* and *UAS Sod2 ^RNAi^* has been reported previously[32, 33]. To confirm the effect of *Sod1* knockdown, *Sod1* mimic line (insertion allele) was utilized, and similar results were obtained with both the RNAi and mimic lines. Since our analysis has been done using the same genotype in different diets, the genetic background of the experimental strain in the different diets is identical. Hence, the statistical difference in lifespan in the experiment can be attributed to the effect of Shankhpushpi versus the solvent-formulated diet. Unless otherwise noted, all lifespan assays utilized adult mated (48h) female flies of the indicated ages. All the experiments were performed at 25°C.

### 2.2 *Drosophila* medium preparation and maintenance

#### 2.2.1. Cornmeal/Agar Food

The cornmeal/agar food was prepared by mixing 86 g of cornmeal, 25g of sucrose, 51g of Dextrose, 15g of Yeast extract, 6g of agar, 1% Acid mix (10 ml) and Tegosept (5ml; 1 gm of Methyl 4-hydroxybenzoate [SRL] in 5ml 100% Ethanol) per 1000 mL of food. The acid mix stock solution was prepared by combining propionic acid (164 ml milliQ water with 836 ml of propionic acid [SRL]) and orthophosphoric acid (917 ml MilliQ water to 83 ml of orthophosphoric acid [SRL]).

#### 2.2.2. Shankhpushpi-supplemented food

Shankhpushpi food was made by dissolving 10 mg shankhpushpi extract in 1 ml of water overnight at 4°C with continuous shaking (For Figures 1, 4, and 6). Shankhpushpi was dissolved in Ethanol for Experiments performed in Figures 2, 3, and 5. Pure Shankhpushpi extract was procured from a readily available commercial source (Nature’s Velvet Lifecare Shankhpushpi pure extract available as 500mg capsules). A 10 mg/ml stock was prepared fresh for each experiment and stored at 4°C, and was diluted to 0.1 mg/ml. To prepare 25X, 50X, and 100X shankhpushpi supplemented food, 181.3µL, 362.9 µL, and 725.8 µL of 0.1mg/ml shankhpushpi extract were added to 50 mL standard yeast-cornmeal food, respectively. Parental generation *Canton S* flies were allowed to lay eggs on shankhpushpi-supplemented medium. The progeny flies were allowed to feed on this food throughout development and behavioural assays were then performed with 2-5 days old adult flies.

#### 2.2.3. Paraquat Food

Methyl viologen dichloride, Paraquat (Sigma Aldrich), was used to study the ability of flies fed on shankhpushpi food to resist the oxidative stress induced. To make a 5mM Paraquat solution, 64 mg of Paraquat was weighed and dissolved in 500μL of water. 50 mL of cornmeal food was melted and cooled down to 40°C. The Paraquat solution was added to the melted food. It was mixed evenly by stirring and layered 2 mL each on 1.5% Agar vials.

#### 2.2.4. Fluoxetine administration

Fluoxetine (Flutax-20; Leeford, India) was diluted to a concentration of 10mM in the *Drosophila* medium [34]. The medium was cooled to 50°C before addition of the powder. The medium was pipetted into isolation vials, and flies were administered the fluoxetine diet for 18 hours.

### 2.3 RNAi-mediated knockdown of target genes

*Actin GS* (Gift from Pankaj Kapahi’s laboratory) virgin flies were crossed to *UAS CG6293^RNAi^* and *UAS Sod1^RNAi^* male flies. Crosses were set on 50x Shankhpushpi-supplemented food. Progeny were allowed to grow on this food. Then, 2-5 days old F_1_ flies were segregated into groups of 10 flies each, and males and females were kept separately. Flies were then randomly assigned to two groups, and one group was fed 50x Shankhpushpi food containing 1 ml RU-486 (4.34 mg/ml) for 5 days, while the other group was fed 50x Shankhpushpi food containing 1 ml ethanol for 5 days. These flies were then subjected to a short variable stress (SVS) regime followed by SPT.

### 2.4 Measurement of food intake

The food intake in flies fed the Shankhpushpi diet was measured by the EX-Q assay[33, 35]. Ex-Q tubes were 14 mL round-bottom dual-position snap cap tubes (Tarsons 860020). The inbuilt cavity in the cap of the tube was used as a food container. Six air holes were made in the cap around the cavity using a pushpin. For Shankhpushpi food used in EX-Q experiments, 1% agarose (Seakem LE agarose, Lonza 50004) was used for preparation instead of agar. The Erioglaucine (SRL 98188, for dye food) was added to the medium after cooling to 60°C.

Three-day old adult mated female or male flies were acclimatized to the experimental food for 48h for EX-Q assays in Figure panels 1A-B. Shankhpushpi-cornmeal medium containing 2.5% (w/v) Erioglaucine was used as the assay medium. Absorbance of solubilized excreta (1000μl) was measured at 630nm (Molecular Devices, SpectraMax i3x) in a 96-well plate. The food intake per fly was calculated from a standard curve prepared from stock solutions of pure dye (0.01-0.065 mg/ml). The absorbance of a 200 µL sample was measured at 630nm. A minimum of 6 replicates of 10 flies each were used for assays, and individual data points were plotted in each panel, and the number of replicates was noted in the figure legends. The blank calculations were performed by preparing homogenates from age-matched flies that were fed a food that lacked Erioglaucine.

### 2.5 Short Variable Stress (SVS) regime for inducing depression-like state in Drosophila

The Short Variable Stress (SVS) regime was adapted from a previously with some modifications [34]. Briefly, 2-5-day-old adult flies were sorted according to sex in empty glass vials, containing 10 flies each[34]. Male flies were exposed to 37°C for 20 minutes, fasted in empty vials for 6 hours, individually housed in 14 mL tubes for 18 hours, and placed in a -5°C ice-water bath for 10 minutes. Female flies were exposed to 37°C for 30 minutes, fasted for 8 hours, socially isolated in individual 14 ml tubes for 18 hours and introduced to a -5°C ice-water bath for 2 rounds of cold shock, for 10 minutes each. Flies were then allowed to completely recover from cold anaesthesia before being used for behavioural assays.

### 2.6. Sucrose Preference Test (SPT)

Control and SVS regime subjected flies were transferred to empty vials in groups of 10 and uniformly fasted for 1.5 hours to allow them to develop an appetite. The cotton plugging of vials were then replaced with sponge cap containing two holes. Two capillaries containing 5 µL each of autoclaved RO water coloured with blue food dye and 5% sucrose solution coloured with red food dye were introduced through these sponge caps. Flies were then allowed to consume both liquids for 3 hours. A control vial without flies was kept to account for evaporative loss. After 3 hours, the capillaries were removed, and the liquid left in the capillary was measured. All vials were kept at 24°C and at 50% RH. Assays were performed during the daytime of their circadian cycle. The Sucrose preference index was calculated using the following formula:

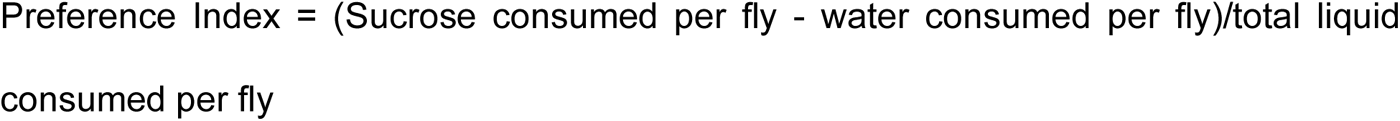

### 2.7. Sample preparation for metabolomic analysis

Samples were prepared from 25 adult flies exposed to a control corn meal diet or Shankhpushpi-supplemented diet. Briefly, the samples were frozen in liquid nitrogen in 2 mL screw cap vials with ceramic beads and stored in -80°C until preparation of the lysate. The sample lysate was prepared by homogenizing the samples in 90% chilled methanol, followed by one hour incubation at -20°C to precipitate proteins, and the sample was centrifuged to remove insoluble debris. The supernatant was dried in a SpeedVac, and the metabolite containing precipitate was analysed.

### 2.8. RNA isolation and Quantitative real-time PCR

Total RNA was extracted from 5-10 adult flies using RNAiso Plus (Takara Bio, Inc)[36, 37]. Animals were homogenized in 0.2 ml of RNAiso Plus with a white micropestle (Tarsons) prior to extraction. The cDNA was generated by using a High-Capacity cDNA Reverse Transcription Kit (Thermo Fisher Scientific, MA, USA). In each reaction 0.5-1μg of RNA was mixed with random hexamers, MgCl_2_, 10X RT Buffer, dNTPs, RNAse Inhibitor and MultiScribe Reverse transcriptase in a 10μL total volume. The cDNA synthesis was performed as per the manufacturer’s protocol in a Bio-Rad C1000 Touch

Thermal Cycler or Bio-Rad T100 Thermal Cycler. The synthesized cDNA was diluted (1:5) and used as template for quantitative real-time PCR (qRT-PCR) using SYBR premix EX-taq plus (TaKaRa) and analyzed on Quant Studio 6 Real-Time PCR machine (Applied Biosystems, Foster City, CA, USA) or Bio-Rad CFX Opus 96 real-time PCR system. The expression of the target genes was normalized to *actin-5C*.

Primers used for RT-PCR are: act-5c For, cacaccaaatcttacaaaatgtgt; act-5c Rev, aatccggccttgcacatg; Sod 2 For, agaacctctcgcccaacaag; Sod 2 Rev, actgcagatagtaggcgtgc; Catalase For, caaccccttcgatgtcacca; Catalase Rev, catcctggttgtccgtcaca; Sod1 For, caagggcacggttttcttcg; Sod1 Rev, tacggattgaagtgcggtcc; HSP70 For, tggacaagtgcaacgacact; HSP70 Rev, tcgatcgaaacattcttatcagtct; Dop1R1 For, tgatacttgggtggcctttg; Dop1R1 Rev, cagagcgcatgttggatact; Rdl For, cgacctggtgtagaaacactatc; Rdl Rev, ccgatgacccaagaccttaaat; GABA-B-R1 For, cgaggagggttacgacattaac; GABA-B-R1 Rev, ggcaatctccttgtccgtatag; CG6293 For, tcgagcacgcagctgaaa; CG6293 Rev, ccgcacagaccccacatt; Glut 1 For, tttcctatgtgcaccgggtc; Glut 1 Rev, gcgaaaatggataccgccac.

### 2.9. Survivability assay

The Kaplan-Meier method was utilized to generate survival curves for all lifespan experiments using the *Online Application for the Survival Analysis of lifespan assays* (OASIS) and GraphPad prism and the p-values were calculated using the log-rank (Mantel-cox) test (Supplementary Table 4) [38, 39]. All experiments were performed with flies that were allowed 48 h to mate after emerging as adults. On the third day after eclosion, flies were anesthetized with carbon dioxide, sorted, and distributed between different dietary regimes for experiments. Around 20 flies were transferred to each vial. For longevity assay live flies were transferred to fresh medium every 2 day and counted every alternate day until no flies remained.

### 2.10. Mass Spectrometry Analysis of Samples

All GC-MS analysis was performed at the Core facility in the university of Utah with an Agilent 5977b GC-MS MSD-HES and an Agilent 7693A automatic liquid sampler. Dried samples were suspended in 40 µL of a 40 mg/mL O-methoxylamine hydrochloride (MOX) (MP Bio #155405) in dry pyridine (EMD Millipore #PX2012-7) and incubated for one hour at 37 °C in a sand bath. 25 µL of this solution was added to auto sampler vials. 60 µL of N-methyl-N-trimethylsilyltrifluoracetamide (MSTFA with 1%TMCS, Thermo #TS48913) was added automatically via the auto sampler and incubated for 30 minutes at 37 °C. After incubation, samples were vortexed and 1 µL of the prepared sample was injected into the gas chromatograph inlet in the split mode with the inlet temperature held at 250°C. A 5:1 split ratio was used for analysis for the majority of metabolites. Any metabolites that saturated the instrument at the 5:1 split were analyzed at a 50:1 split ratio. The gas chromatograph had an initial temperature of 60°C for one minute followed by a 10°C/min ramp to 325°C and a hold time of 10 minutes. A 30-meter Agilent Zorbax DB-5MS with 10 m Duraguard capillary column was employed for chromatographic separation. Helium was used as the carrier gas at a rate of 1 mL/min.

Data was collected using MassHunter software (Agilent). Metabolites were identified and their peak area was recorded using MassHunter Quant. This data was transferred to an Excel spread sheet (Microsoft, Redmond WA). Metabolite identity was established using a combination of an in-house metabolite library developed using pure purchased standards, the NIST library and the Fiehn library. Data was analyzed using in-house software to prepare for analysis by the “MetaboAnalyst” software tool.

### 2.11. Statistical analysis

Statistical analysis and data presentation were performed with GraphPad Prism 10 software, OASIS, and Microsoft Excel. Survival curves were compared using log-rank tests and in figures where multiple comparisons were made p-values were calculated after applying Bonferroni’s correction and noted in the figure legends and Tables. All survival and lifespan graphs show two repeats with cohorts of 150-200 female flies per genotype with 20 files per vial. An ordinary one-way ANOVA was used to analyze data from fluoxetine treatment where multiple comparisons were made. All RT-PCR analysis was performed with three independent biological replicates and two technical replicates for each biological replicate and individual data points were plotted in all graphs. The Fisher Least Significant Difference (LSD) Method was used to compare means from multiple processes and adjusted p-values for all comparisons were computed by applying Bonferroni’s correction and noted in the figure legends. The significant p-values were denoted in black font and the non-significant values were noted in red font. Statistical significance was set at p<0.05.

Metabolomic datasets were analysed using MetaboAnalyst 5.0. Data was analysed using one-factor statistical analysis. Raw peak intensities were entered in rows. Data was then normalized according to median and autoscaling was performed. Variance filter was set at 5% Mean Absolute Deviation (MAD) and Abundance filter was set at 0% Median Intensity Value. Principal Component Analysis (PCA) was performed, and the scores plot was obtained. The volcano plot was generated using the same parameters.

## 3. RESULTS

### 3.1 Short variable stress induces anhedonia in *Drosophila melanogaster*

To establish a model for assessing the psychopharmacological properties of *C. pluricaulis* (Shankhpushpi), we optimized a protocol for inducing stress in both male and female adult flies. This protocol combined psychosocial and physical stressors to develop anhedonia-like behavior, resembling a depression-like state. 3-5-day-old flies were sorted according to sex and subjected to a stress regime consisting of heat shock at 37°C, fasting, social isolation, and cold shock at -5°C (**Figure 1A**). Following exposure to the stress regime, unstressed control flies and stressed flies were offered a choice between 5% sucrose and water, and their anhedonia was tested through the sucrose preference test (SPT) for 3 hours. Anhedonia is the lack of interest in enjoyment or pleasure, which is a typical symptom of a depression-like state [40]. Wild-type male and female flies respond to the stress regime by showing reduced preference for sucrose (Preference Index {PI} dropped from 0.4754 ± 0.3343 to -0.2812 ± 0.4080 in males and from 0.5427 ± 0.2739 to -0.4668 ± 0.3593 in females), indicating disinterest in reward-seeking activities (Figure 1B-C; Panels 1 and 3). Selective Serotonin Reuptake Inhibitors (SSRIs) are known to reverse depression-like behaviours by binding to pre-synaptic neurons and enhancing serotonin signalling in the brain. Fluoxetine is one such SSRI known to relieve depression like behaviour in mice and humans, and we wanted to assess its effects on behaviour in response to our stress regime in flies. To ensure that this behavior was reversible and a response to the stress regime alone, we administered 10 mM fluoxetine to stressed flies and tested their behavior in SPT. Fluoxetine administration led to the reversal of anhedonia in stressed male and female *Canton S* flies (the PI index in treated flies increased from -0.1492 ± 0.2055 to +0.4406 ± 0.1097 in males and from -0.05587 ± 0.3674 to 0.8679 ± 0.09840 in females upon exposure to Fluoxetine). Their preference levels recapitulated the values obtained for unstressed flies (**Figure 1B-C, panels 2 and 4)**. Thus, we found that administering 10 mM fluoxetine for 18 hours during the social isolation period led to relief from depression-like behavior in wild-type flies, while control flies showed no significant alterations in behavior upon fluoxetine treatment (**Figure 1B-C)**. This amelioration of the variable stress-induced phenotype suggests that evolutionarily conserved biochemical pathways are involved in the depressive state of flies and Major depressive disorder (MDD) in humans, and that this model could be utilized for studying the effects of other pharmacological interventions and the underlying conserved genetic players.

### 3.2. Dietary administration of *C. pluricaulis* alleviates stress-induced anhedonia in Drosophila

To understand the effect of dietary intake of *C. pluricaulis* (Shankhpushpi) on stress-responsive behavior, we administered two different dosages of Shankhpushpi-supplemented regular fly food to wild-type *Canton S* flies throughout the developmental period and adulthood. 3-5-day-old flies were sorted according to sex and subjected to the short variable stress regime (**Figure 1D**). To ensure this effect was due to administration of Shankhpushpi alone and not due to altered food consumption upon Shankhpushpi supplementation, we measured the food intake of flies by performing the Excreta-Quantification (Ex-Q) assay to measure food intake per fly. We found supplementation of 50X Shankhpushpi in fly food did not affect food consumption of *Canton* S male and female flies (**Figure 1E-F**).

Following exposure to the stress regime, control flies raised on a standard fly diet and Shankhpushpi-fed flies were offered a choice between 5% sucrose and water. Their anhedonia-like behavior was tested through the sucrose preference test (SPT) (**Figure 1G-H**). Dosage was calculated according to the recommended intake prescribed to human beings, standardized to approximate the body weight of flies. We found that dietary supplementation of 25X Shankhpuhspi extract in regular fly food resulted in alleviation of anhedonia in male *Canton S* flies. At the same time, no significant effect was observed in female flies **(Figure S1C-D)**. We further increased the dosage of Shankhpushpi to 50X and found that both male and female *Canton S* flies raised on 50X Shankhpushpi-supplemented food showed relief from anhedonia, as they preferred more sucrose than water, compared to their control counterparts raised on regular fly food and exposed to stress. These control flies lost their preference for sucrose upon exposure to a psychosocial and physical stress regime, indicating an anhedonia-like state. At the same time, Shankhpushpi administration alleviated this stress-induced anhedonia-like behavior (PI increased from -0.3257 ± 0.1341 to +0.3663 ± 0.2936 in males and from - 0.1801 ± to 0.1199 ± 0.03973 in females that were administered Sh diet) (**Figure 1G-H**). These results indicate that Shankhpushpi has anti-depressant properties when administered at 50X concentration in wild-type flies.

**Figure 1.**
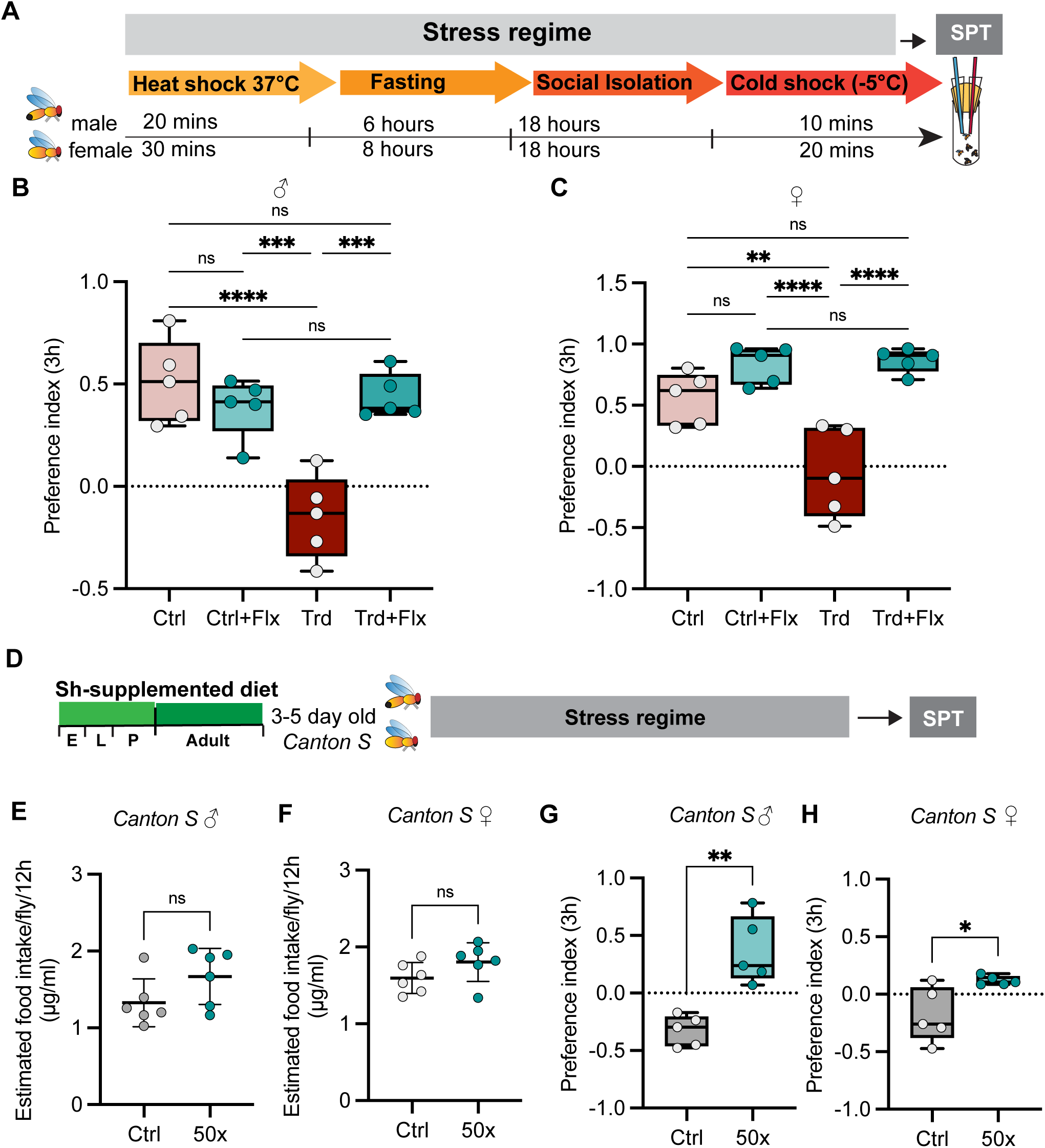
Administration of *Convolvulus pluricaulis* ameliorates stress-induced anhedonia in wild-type flies. (A) A schematic diagram representing the stress regime for inducing a depression-like state in *Drosophila melanogaster*. 3-5 Day old flies were exposed to a stress regime comprising of a Heat shock at 37°C (20 minutes for males and 30 minutes for females), fasting (6 hours for males and 8 hours for females), 18h of social isolation and cold shock at -5°C (10 minutes for males and 20 minutes for females) following which sucrose preference test was performed for 3h. (B and C) The preference index of male *Canton S* flies (B) and female *Canton S* flies (C). Each circle represents the preference index using 10 flies. The box plot shows the median and the box spans from the first quartile to the third quartile. The whiskers extend from the box to the minimum and maximum data points. Panel 1 represents Control (Ctrl) flies that are not exposed to stress, the Panel 2 represent control flies treated with 10mM Fluoxetine (Ctrl +Flx), the third panel represents flies that were exposed to a stress regime, Treated (Trd) and the fourth panel represents the flies that were treated with 10mM Fluoxetine followed by exposure to stress regime (Trd +Flx). Statistical significance was determined using a one-way ANOVA. Adjusted P-values after applying Bonferroni’s correction are represented in the panel, and we used an α level of 0.05 to assess statistical significance. (D) The experimental scheme utilized to determine the impact of *C. pluricaulis* in the stress-induced depression model. *Canton S* flies were fed Shankhpushpi throughout development and adulthood (E, embryonic stage; L, larval stage; A, adult stage). 3-5-day-old flies were subjected to a stress regime, and their anhedonia-like behavior was measured using the sucrose preference test (SPT). (E-F) Male (E) and female (F) *Canton S* flies consume comparable amounts of normal fly food and 50X Shankhpushpi supplemented food. Food intake was measured by Ex-Q assay. Each data point represents a group of 10 flies. Data are represented as mean ± SD, n=6. P-value was calculated using an Unpaired t-test with Welch’s correction. (G-H) 50X Shankhpushpi-fed male (G) and female (H) *Canton S* flies exposed to stress regime, showing positive preference index, indicating more sucrose consumption and decreased anhedonia in comparison to stress-exposed control flies raised on regular fly food, showing lower preference index indicating anhedonia. Each data point for sucrose preference test (SPT) represents a group of 10 flies. The Box plot shows the median and the box spans from the first quartile to the third quartile. The whiskers extend from the box to the minimum and maximum data points, and we used an α level of 0.05 to assess statistical significance. *p<0.05, ***p<0.005 by Student t-test.

### 3.3. Changes in the metabolome upon intake of Shankhpushpi

To determine how *Convolvulus pluricaulis* intake impacts metabolism in the brain and the rest of the body, we performed small-molecule gas chromatography/mass spectrometry (GC/MS) analysis with head and decapitated body tissue of wild type *Canton S* flies that were fed a control diet or a diet supplemented with *C.pluricaulis* (Shankhpushpi) for 20 days (**Figure 2 and Figure S2 and S3)**. This metabolomic approach revealed that the intake of Shankhpushpi led to a significant increase in L-Ascorbate, Adenosine monophosphate, Myristic acid, and D-Glucose in the head tissue (**Figure 2A, D-G)**. Principal component analysis revealed that the metabolomes of the head tissue show no overlap between their confidence intervals (**Figure 2B**). To determine the factors driving the metabolomic differences between the control and Shankhpushpi-fed flies, we used correlation analysis to identify metabolites altered in response to elevated L-Ascorbate **(Figure S2)**. This approach revealed that L-Ascorbate levels positively correlate with the abundance of D-Glucose, L-Sorbose, Ethanolamine, Adenosine monophosphate and Myristic acid. In contrast, concentrations of Fumaric acid, Succinic acid and D-Malic acid as well as amino acids such as L-Proline and L-Aspartic acid displayed inversely proportional relationships with L-Ascorbate **(Figure S2)**.

**Figure 2.**
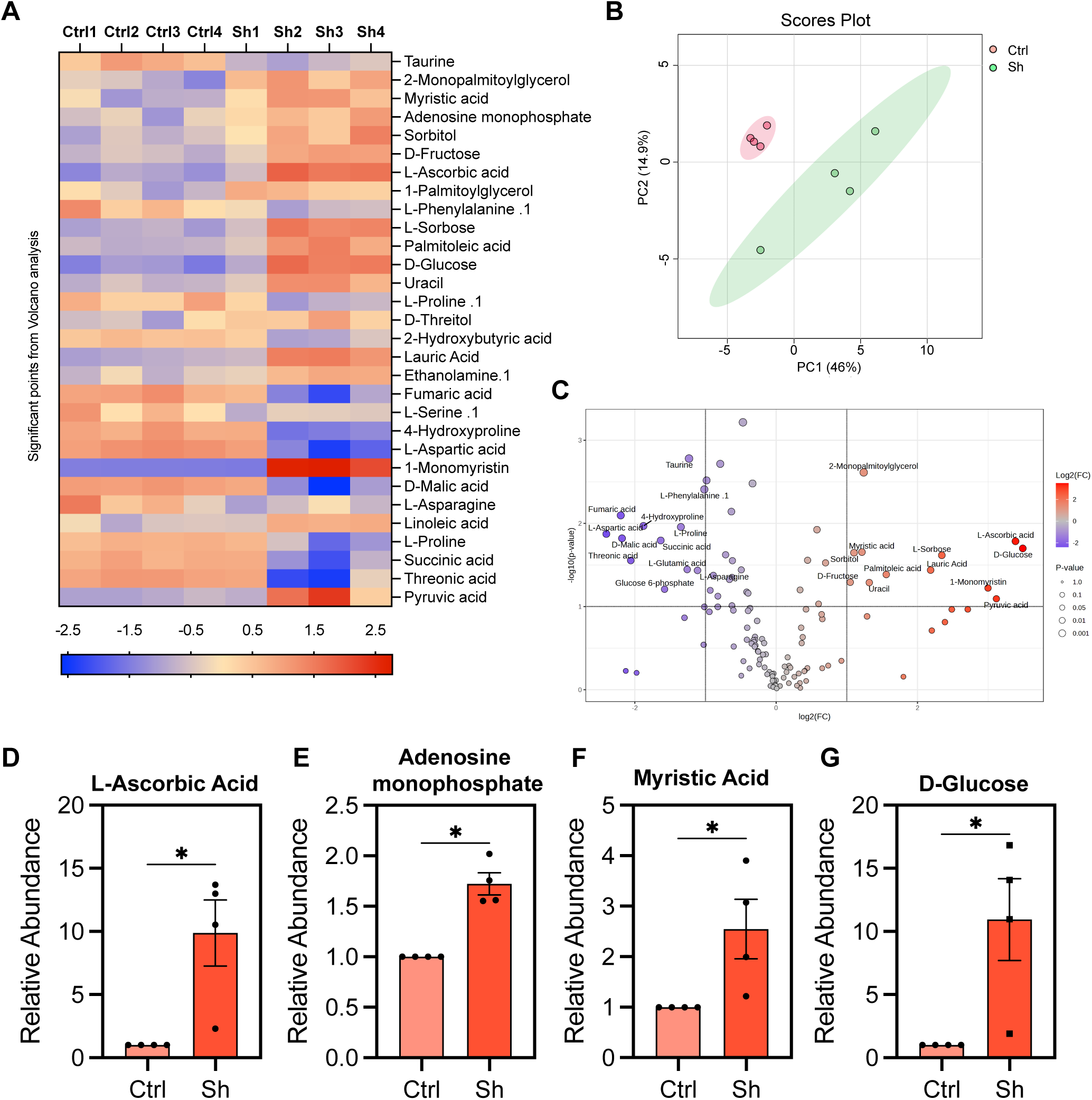
Metabolomic analysis of the head tissue extract of *C. pluricaulis* (Shankhpushpi)-fed wild-type flies. Canton S flies (females) were fed a Control diet or a diet supplemented with Shankhpushpi for 20 days. The head tissue was dissected and analyzed using GC-MS. (A) Heatmap showing the Top 30 significantly altered metabolites in the head tissue of Shanlkhpushpi-fed (Sh) wild-type flies in comparison to flies raised on control diet (Ctrl). Metabolomics analysis was performed using four biological replicates for both the control and Shankhpushpi-fed groups. (B) Projection of Shankhpushpi and control metabolite samples onto principal components 1 and 2 (PC1 vs PC2), explaining 46% and 14.29% of variance, respectively. Red points indicate regular fly food-fed controls, and green points indicate metabolites in Shankhpushpi-fed flies. (C) Volcano plot showing significantly upregulated and downregulated metabolites in Shankhpushpi-fed flies compared to control flies. (D-G) L-ascorbic acid, adenosine monophosphate, myristic acid and D-glucose levels are significantly increased in head tissue of Shankhpushpi-fed flies in comparison to controls. Error bars indicate mean ± S.D. *p<0.05 by Student t-test.

In contrast, no significant changes were observed in the metabolome of the decapitated body tissue of flies that were fed a Shankhpushpi-supplemented diet for 20 days **(Figure S3A-D)**. The Principal Component analysis (PCA) revealed that the metabolome of the decapitated body tissue of flies that were administered Shankhpushpi-diet was similar to the metabolome of flies that were fed a Control diet lacking Shankhpushpi **(Figure S3E)**. D-fructose, Glyceraldehyde-3-Phosphate, and a plant-derived metabolite, Eicosane, were found to be upregulated in the Shankhpushpi group, and were not investigated further. These data were consistent with the previous studies that have confirmed the neuropharmacological properties of this extract. The effect of L-Ascorbate supplementation has been tested in several studies, which have shown that it can function both as an antioxidant and a neuromodulator for the treatment of depression[41]. We predicted that at least some of the beneficial effects were mediated by the increase in L-Ascorbate levels. Hence, we examined whether the expression of genes that facilitate L-Ascorbate accumulation in the brain was modulated by *C. pluricaulis* (Shankhpushpi) intake.

### 3.4. Effect of *C. pluricaulis* (Shankhpushpi) diet on gene expression

*C. pluricaulis* intake leads to a significant increase in L-Ascorbate and Glucose in the brain tissue. Hence, we examined the expression of genes associated with increased expression and transport of L-Ascorbate (*CG6293, GABA-B-R1* and *Rdl*), Glucose (Glut1), and genes involved in the antioxidant effects (*Sod1*) in the brain (**Figure 3**)[42]. We predicted that if some of the beneficial effects of Shankhpushpi were mediated by conserved metabolic pathways, then changes in the expression of genes regulating these pathways would be observed upon intake of Shankhpushpi compared to the Control diet. Total RNA was extracted from the head and body (without Head) tissue of wild-type (*Canton S*) flies that were fed a Control or Shankhpushpi-supplemented diet for 10 days, and RT-PCR was performed to quantify expression of *CG6293*, *glut 1*, *sod1*, *rdl,* and *GABA-B-R1* in the head tissue. The gene expression analysis in the head tissue was performed by feeding three different concentrations of Shankhpushpi (10X, 25X and 100X), while only the highest concentration was tested for the body tissue (100X). Wild type flies that were fed a Shankhpushpi formulated diet for 10 days expressed significantly higher levels of *CG6293* in the head tissue at all three concentrations tested (Sh vs Ctrl [10X]: 2.34 ± 1-fold increase; Sh vs Ctrl [25X]: 1.62 ± 0.35-fold increase; Sh vs Ctrl [100X]: 1.74 ± 0.49-fold increase) (**Figure 3A, F, K)**. The *Drosophila* gene *CG6293* has been predicted to be involved in the transmembrane transport of L-Ascorbic acid. It is orthologous to *SLC23A1*(solute carrier family 23 member 1) and *SLC23A2* (solute carrier family 23 member) in humans and code for two isoforms of the sodium-dependent vitamin C transporters (SVCT1 and SVCT2), respectively. SVCT 1 is the predominant carrier of ascorbic acid in the intestine, and SVCT2 is primarily responsible for transporting Ascorbic acid in the brain[43]. We also examined the expression of two other genes, *GABA-B-R1* (Sh vs Ctrl [10X]: 1.43 ± 0.56-fold increase; Sh vs Ctrl [25X]: 1.65 ± 0.35-fold increase; Sh vs Ctrl [100X]: 1.25 ± 0.12-fold increase) and *rdl* (Sh vs Ctrl [10X]: 1.49 ± 0.43-fold increase; Sh vs Ctrl [25X]: 1.38 ± 0.29-fold increase; Sh vs Ctrl [100X]: 1.67 ± 0.38-fold increase) that have been linked to Ascorbate homeostasis and both were upregulated upon intake of Shankhpushpi were *GABA-B-RI* (Figure 3C, D, H, I, M and N). While the levels of *rdl* increased at all three concentrations of Shankhpushpi, GABA-B-R1 levels were significantly increased at higher concentrations only (25X and 100X). Previous metabolomic studies conducted in humans have linked Ascorbate to GABAergic signaling in depression-like states[44]. Ascorbate has been shown to enhance the function of GABA receptors in the retina[45]. *Drosophila Resistant to dieldrin* (*Rdl)* is the subunit of the GABA_A_ receptor and has been implicated in modulating olfactory learning and regulating sleep [46]. GABA-BR-1 is a receptor known to play an essential role in olfactory perception in *Drosophila*. Consistent with previous studies in *Drosophila* that have linked Ascorbate intake to increased *GABA-B-R1* expression, we found that intake of Shankhpushpi for 10 days led to a significant but moderate increase in expression of both GABA-BR-1 and *rdl* in the head tissue. Next, we examined the expression of *glut1* in the head tissue of Shankhpushpi-fed flies. *Drosophila Glut 1* is a glucose transporter that plays a role in glucose uptake and the storage of triglycerides [47, 48]. A significant increase in expression of *Glut1* was observed at all three concentrations of Shankhpushpi (Sh vs Ctrl [10X]: 1.55 ± 0.5-fold increase; Sh vs Ctrl [25X]: 1.28 ± 0.17-fold increase; Sh vs Ctrl [100X]: 1.28 ± 0.15-fold increase) (**Figure 3B, 3G, 3L**). Finally, we examined the expression of *Superoxide dismutase 1* (*Sod1)*, a gene that plays a conserved critical role in antioxidant defense [49–53]. Reactive oxygen species and reactive nitrogen species are overproduced upon oxidative stress and these reactive species cause oxidation of proteins, nucleic acids, and fatty acids[54]. Sod1 is a cytoplasmic copper/zinc-containing enzyme that catalyzes the conversion of superoxide radicals into hydrogen peroxide[55]. The Catalase enzyme converts hydrogen peroxide into water and oxygen and effectively removes the reactive oxygen species[56, 57]. The levels of *sod1* in the head tissue was found to be increased at all three concentrations of Shankhpushpi (Sh vs Ctrl [10X]: 1.2 ± 0.05-fold increase; Sh vs Ctrl [25X]: 1.2 ± 0.11-fold increase; Sh vs Ctrl [100X]: 1.16 ± 0.12-fold increase) (**Figure 3E, 3J, 3O)**.

**Figure 3.**
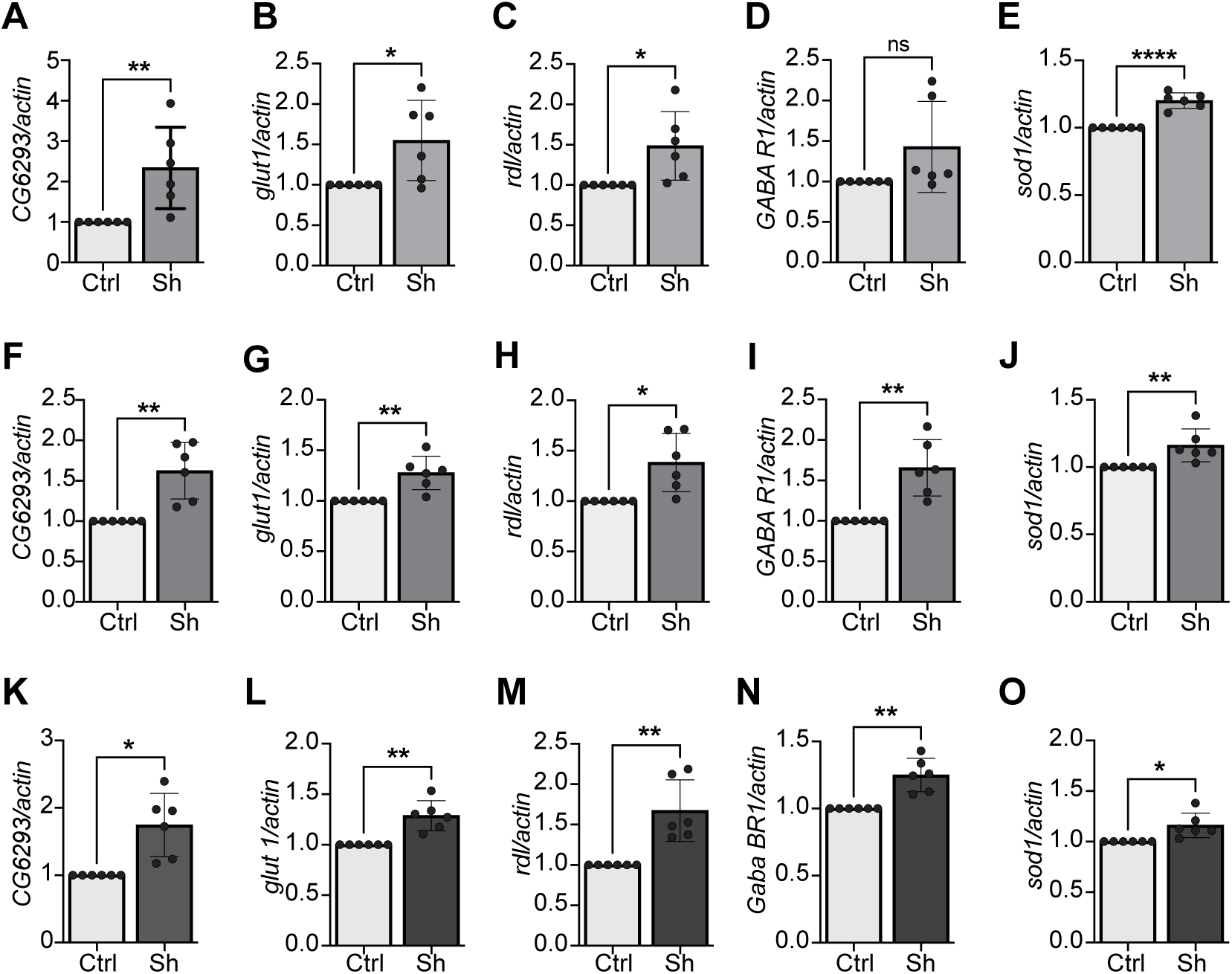
Effect of *Convolvulus pluricaulis* (Shankhpushpi)-formulated diet on expression of genes in the head tissue of wild-type *Drosophila*. Real-time reverse transcriptase-polymerase chain reaction (RT-PCR) quantitation of *CG6293*, *glut1*, *rdl*, *GABA-B-R1* and *sod1* mRNA in head tissue of female *Canton S* flies that were fed a control diet (Ctrl) or a Shankhpushpi (Sh) diet for 10 days. Three different concentrations of Shankhpushpi were administered: 10X (A-E), 25X (F-J) and 100X (K-O). Expression levels were normalized to *actin*. Values are mean ± SD, n= 3. For each biological replicate (n=3), two technical replicates were analyzed. (A-E) At the lowest concentrations (10X), Shankhpushpi significantly increased expression of *CG6293*, *glut1*, *rdl*, *GABA-B-R1* and *sod1 CG6293, glut1, rdl, and sod1* but no significant modulation was observed for *GABA-B-R1*. (F-O) At moderate (25X) and high (100X) concentrations, a considerable upregulation of *CG6293*, *glut1*, *rdl*, *GABA-B-R1,* and *sod1* mRNA expression was observed in the head tissue of wild-type flies. Error bars represent mean ± SD, and p-value was calculated using unpaired t-test with Welch’s correction and is noted in the bar graph. *p<0.05, **p<0.01 and ****p<0.0005 by Student t-test.

To infer whether the antioxidant property of Shankhpushpi was due to changes in gene expression of antioxidant enzymes in tissues other than the head, we examined the expression of ubiquitously expressed genes, including *sod1*, *sod2*, *caspase,* and *hsp70,* in the body tissue (w/o head). Our analysis showed that *sod1* levels were increased and *sod2* levels were significantly decreased upon intake of Shankhpushpi (Sh vs Ctrl [100x]: 0.86 ± 0.07-fold decrease); *sod1* (Sh vs Ctrl [100x]: 1.4 ± 0.26-fold increase) (**Figure 4A-B**). However, no significant change was detected for *catalase* and *hsp70*, indicating that the beneficial effects of Shankhpushpi were likely not mediated by *catalase*, and *hsp70* (**Figure 4C-D**). In summary, our candidate gene expression analysis indicates that Shankhpushpi modulates the expression of conserved genes involved in ascorbate homeostasis, glucose transport, and defense against oxygen radicals.

**Figure 4.**
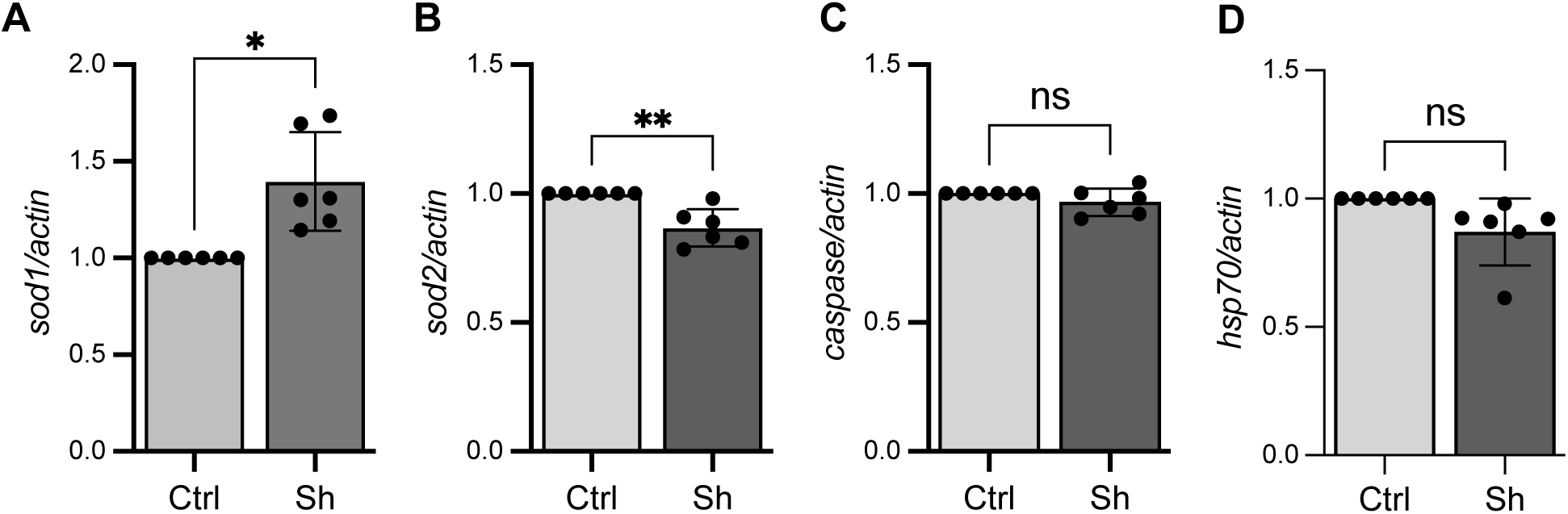
*Sod1* and *Sod2* levels are significantly altered in the body tissue of *Convolvulus pluricaulis* (Shankhpushpi)-fed flies. Real-time reverse transcriptase-polymerase chain reaction (RT-PCR) quantitation of sod1, sod2, caspase, and hsp70 mRNA in the body tissue of female *Canton S* flies that were fed a control diet (Ctrl) or a 100X Shankhpushpi (Sh) diet for 10 days. (A) *Sod1* levels are upregulated in the body tissue of Shankhpushpi-fed flies in comparison to solvent-fed controls. (B) *Sod2* is significantly downregulated in the body tissue of Shankhpushpi-fed flies in comparison to their controls. (C-D) *Caspase* and *hsp70* levels remain unchanged in the body tissue of Shankhpushpi-fed flies in comparison to controls. Error bars represent mean ± SD, and p-value was calculated using unpaired t-test with Welch’s correction and is noted in the bar graph. *p<0.05, **p<0.01 and ^ns^p>0.05 by Student t-test.

### 3.5. *CG6293* is required for the antidepressant effect of Shankhpushpi (Sh)

Metabolomic analysis of head tissue extract of Sh-fed flies revealed several metabolites, such as L-ascorbate, to be upregulated in Sh-supplemented samples (**Figure 2D**). Gene expression analysis also revealed that *CG6293*, L-ascorbate transporter in *Drosophila*, is upregulated in head and body tissue of flies raised on Shankhpushpi diet (**Figure 3 A, F, K)** along with other genes like *Sod1*, *Rdl* and *GABA*. To delineate the molecular effectors through which the anti-depressant effect of Shankhpushpi is mediated, we knocked down *CG6293* and *Sod1* ubiquitously in adult flies by crossing the Actin Gene-switch Gal4 driver with UAS RNAi lines. F_1_ progeny were then sorted according to sex and administered RU-486 containing food to ensure knockdown of relevant genes, along with dietary supplementation of Shankhpushpi (**Figure 5A**). Experimental controls were administered food containing ethanol and Shankhpushpi extract. Knockdown of the functionally relevant mediator of anti-depressant effect of Shankhpushpi would abolish the beneficial effect of Shankhpushpi even in its presence. Knockdown of *CG6293* and *Sod1* was confirmed by qRT-PCR for the respective genes (**Figure 5 B-C, F-G)**. To test for anhedonia-like behavior, we exposed knockdown and control flies to a stress regimen and subjected them to a sucrose preference test. Subsequently, we found that knockdown of L-ascorbate transporter, *CG6293,* throughout the body led to exacerbation of the antidepressant effect of dietary administration of 50X Shankhpushpi (PI decreased from 0.2900 ± 0.1252 to -0.01400 ± 0.1649 in males and from 0.1749 ± 0.1776 to -0.3351 ± 0.1878 in female flies) (**Figure 5D-E**) in both male and female flies. However, knockdown of *Sod1* had no such adverse effect on preference index in the presence of Shankhpushpi in both male and female flies and no statistical significant difference was observed in the PI of the control or RNAi line (Control (male) PI: 0.03449 ± 0.1713; *Sod1^RNAi^* (male): 0.1475 ± 0.1181; Control (female) PI: 0.2525 ± 0.3218; *Sod1^RNAi^* (female): 0.006931 ± 0.1134) (**Figure 5H-I**), indicating it may not be involved in the regulation of behavioral effects of Shankhpushpi. Our data indicate that Shankhpushpi imparts stress-resilience upon its dietary intake through the L-ascorbate transporter (*CG6293*) in both male and female *Drosophila*.

**Figure 5.**
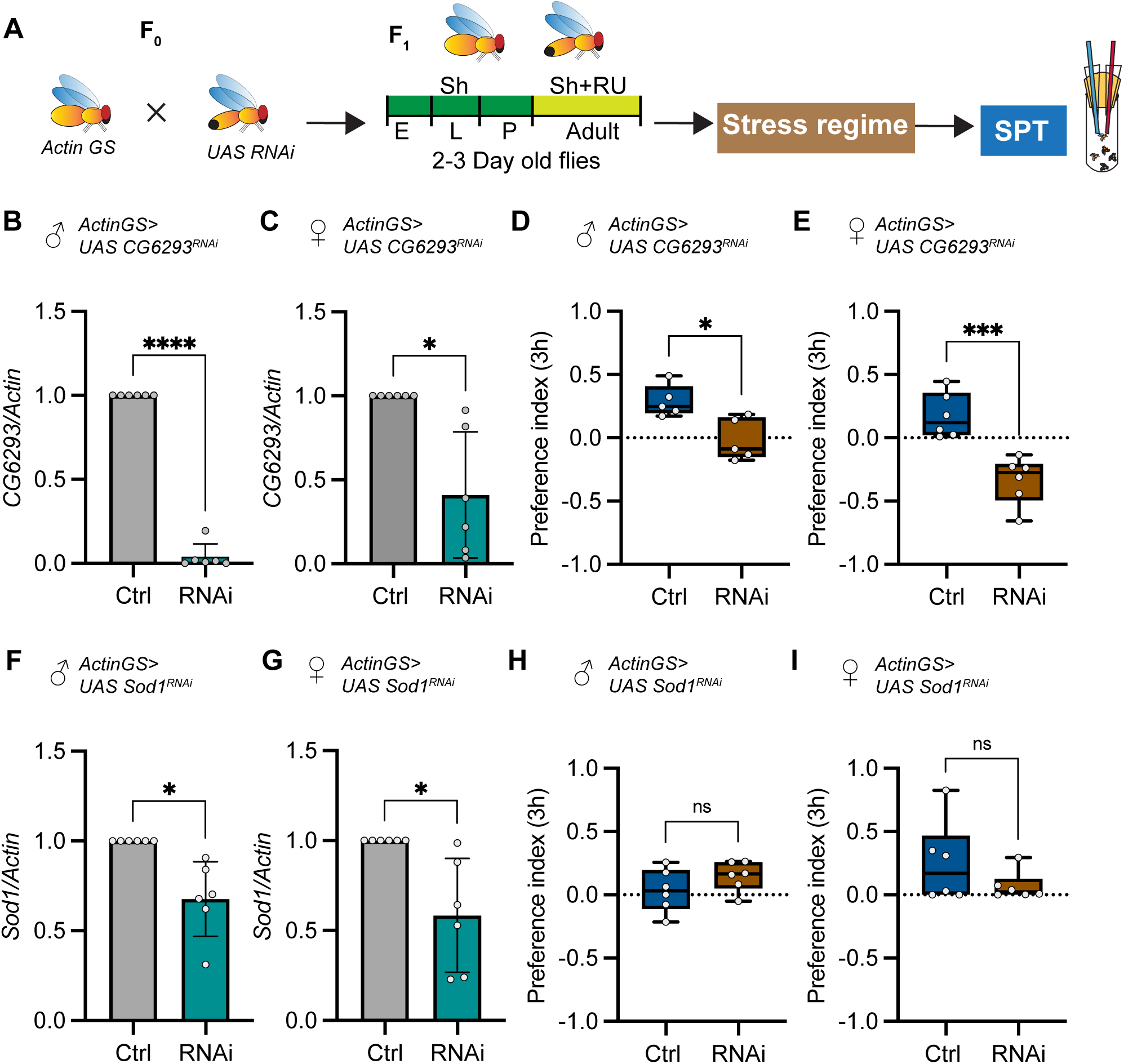
Antidepressant effect of *Convolvulus pluricaulis* (Shankhpushpi) supplementation is mediated by the Ascorbate transporter (*CG6293*). (A) Dietary scheme for analysis of *ActinGS>UASCG6293^RNAi^* and *ActinGS>UASCGSod1^RNAi^* flies. *ActinGS>UAS^RNAi^* flies were raised on 50X Shankhpushpi-supplemented diet (E, embryonic stage; L, larval stage; A, adult stage). Two-to three-day-old adult flies were then fed a 50X Shankhpushpi diet containing RU-486 or solvent (control) for 5 days to knock down the expression of the gene of interest throughout their entire body. Flies were then subjected to a stress regime, and their anhedonia-like behavior was assayed by the sucrose preference test (SPT). (B-C) *CG6293* knockdown in whole body extract of male (B) and female (C) flies fed RU-486, normalized to solvent-fed controls. RNA was extracted from the whole body of 4-5 flies per biological replicate. (D-E) *ActinGS>CG6293^RNAi^* male (D) and female (E) stressed flies fed RU-486 in the presence of 50X Shankhpushpi showing decreased preference index indicating anhedonia in comparison to control flies fed 50X Shankhpushpi and solvent showing positive sucrose preference and resilience to stress.(F-G) *Sod 1* knockdown in whole body extract of male (F) and female (G) flies fed RU-486 normalized to ethanol fed controls. RNA was extracted from the whole body of 4-5 flies per biological replicate. (H-I) *ActinGS>Sod1^RNAi^* male (H) and female (I) stressed flies fed RU-486 in the presence of 50X Shankhpushpi showed no significant difference in sucrose preference in comparison to control stressed flies that were fed 50X Shankhpushpi and solvent, showing resilience to stress-induced anhedonia. (D-E and H-I) Each data point for sucrose preference test (SPT) represents a group of 10 flies. The box plot displays the median, with the box spanning from the first quartile to the third quartile. The whiskers extend from the box to the minimum and maximum data points, and we used an α level of 0.05 to assess statistical significance. (B-I) Error bars represent mean ± SD, and p-value was calculated using unpaired t-test with Welch’s correction and is noted in the bar graph. *p<0.05, **p<0.01 and ^ns^p>0.05 by Student t-test.

### 3.6. The antioxidant activity of Shankhpushpi (Sh) is conferred by the activity of *sod1*, *sod2*, *glut1*, *rdl*, *GABA-B-R1*, and *CG6293*

Shankhpushpi exhibits antioxidant properties in both in vitro and in vivo assays. It has been proposed to function by scavenging free radicals in the 2,2-diphenyl-1 picrylhydrazyl assay (DPPH assay) [20, 58]. In animal models and neuronal cell lines, Sh has been shown to provide neuroprotective effects [59, 60]. Though several phytochemical constituents of Sh have been linked to its neuropharmacological effects, the precise mechanisms underlying these activities remain unknown. Here, we utilized *Drosophila* as a model to unravel the genes responsible for the antioxidant property of Sh. Paraquat (PQ) is a compound used to induce oxidative stress in various animal models, including *Drosophila melanogaster* [61]. Its administration leads to the production of highly reactive superoxide anions, hydrogen peroxide, and hydroxyl radicals, which damage cellular macromolecules and oxidize reducing agents like NADPH and reduced glutathione, essential for normal cell function [62]. Our previous study, along with others, has demonstrated that PQ-induced oxidative stress can be used to evaluate the effectiveness of antioxidants in *Drosophila melanogaster* [33, 63]. To identify downstream effectors mediating the antioxidant effect of Shankhpushpi, the survivability of stressed flies was measured with or without Shankhpushpi (**Figure 6A**). These experiments were performed using strains in which the expression of genes modulated by Shankhpushpi intake was reduced genetically or via RNA interference (**Figure 6 B-O**). We hypothesized that knocking down functionally relevant downstream effectors would increase sensitivity to the toxic effects of PQ, even in the presence of Sh. Therefore, we anticipated that reducing the levels of these key genes would eliminate the lifespan extension effect of Shankhpushpi under PQ treatment. Since Shankhpushpi influences the expression of *rdl*, *sod 1*, *sod 2*, *GABA-B-R 1*, *glut 1*, and *CG 6293*, we examined strains with decreased expression of each of these genes. 2-day-old flies with loss of one copy of *rdl*, *sod 1*, *GABA-B-R 1*, *glut 1*, and *CG 6293* were transferred to food containing either solvent or Shankhpushpi in 5 mM PQ. To compare the effects of a Shankhpushpi-supplemented diet in *Sod1-* and *Sod2*-deficient flies, *sod1* and *sod2* knockdowns were induced ubiquitously in adult flies using a steroid (RU-486) inducible system with the *Daughterless* GeneSwitch Gal 4 (*daGS*) driver (**Figure 6 D-G**). RT-PCR analysis of total RNA extracted from whole animals of *daGS/UAS sod 2 ^RNAi^* and *daGS/UAS sod 1 ^RNAi^* female flies confirmed the knockdown of *sod1* and *sod2* **(Figure S4 C-D)**. Induction of *UAS sod1 ^RNAi^* and *UAS sod2 ^RNAi^* resulted in approximately 32 ± 10.7% reduction in *sod1* levels and approximately 56.6 ± 9.9% reduction in *sod2* levels upon treatment with RU-486 for 5 days in female flies **(Figure S4 C-D)**. *daGS/UAS sod1 ^RNAi^*female flies fed a Shankhpushpi diet had a 10.5% decrease in median lifespan in both experimental trial 1 and experimental trial 2 (**Figure 6F-G, and Table S3**). However, knockdown of *Sod2* did not lead to a reduction in median or maximum lifespan. The Shankhpushpi-fed flies continued to have a 22% increase in median lifespan and a 14.2% and 28% increase in maximum lifespan in Experiments 1 and 2, respectively (**Figure 6D-E, Table S3**). Genetically reducing one copy of *sod1, GABA-B-R1*, *glut1* and *CG6293* did not lead to any increase in survivability in Sh-fed flies under Paraquat-induced oxidative stress and reduced dosage of *rdl* led to a 18% and 8% decrease in median lifespan and in Experiment 1 and 2, respectively (**Figure 6F-O; Table S3)**. *Sod1* codes for the antioxidant enzyme, Superoxide dismutase 1, which is an evolutionarily conserved Reactive oxygen species (ROS) neutralizing enzyme that converts superoxide anions to Hydrogen peroxide [64]. In *Drosophila*, *Rdl* (a GABA-A receptor subunit) and *GABA-B-R1* play crucial roles in gamma-aminobutyric acid (GABA) signaling, and GABA shunt plays a role in maintaining redox balance in blood progenitor cells[65]. The orthologs of glucose transporter (*Glut1*) and ascorbate transporters (*CG6293*) have been linked to oxidative stress defense in other models. Glut1 is a transporter for both Glucose and dehydroascorbic acid, and mutations in this gene or inhibitors that prevent glucose transport have been shown to increase ROS levels in myoblasts[66]. Though *CG6293* is not well characterized in *Drosophila*, it is an ortholog of Ascorbate transporters SVCT1 and SVCT2 and L-Ascorbate or Vitamin C has been shown to function as an antioxidant in several biological systems [67]. Taken together, these analyses have uncovered previously unknown, conserved molecular effectors responsible for the antioxidant property of *Convolvulus pluricaulis*.

**Figure 6.**
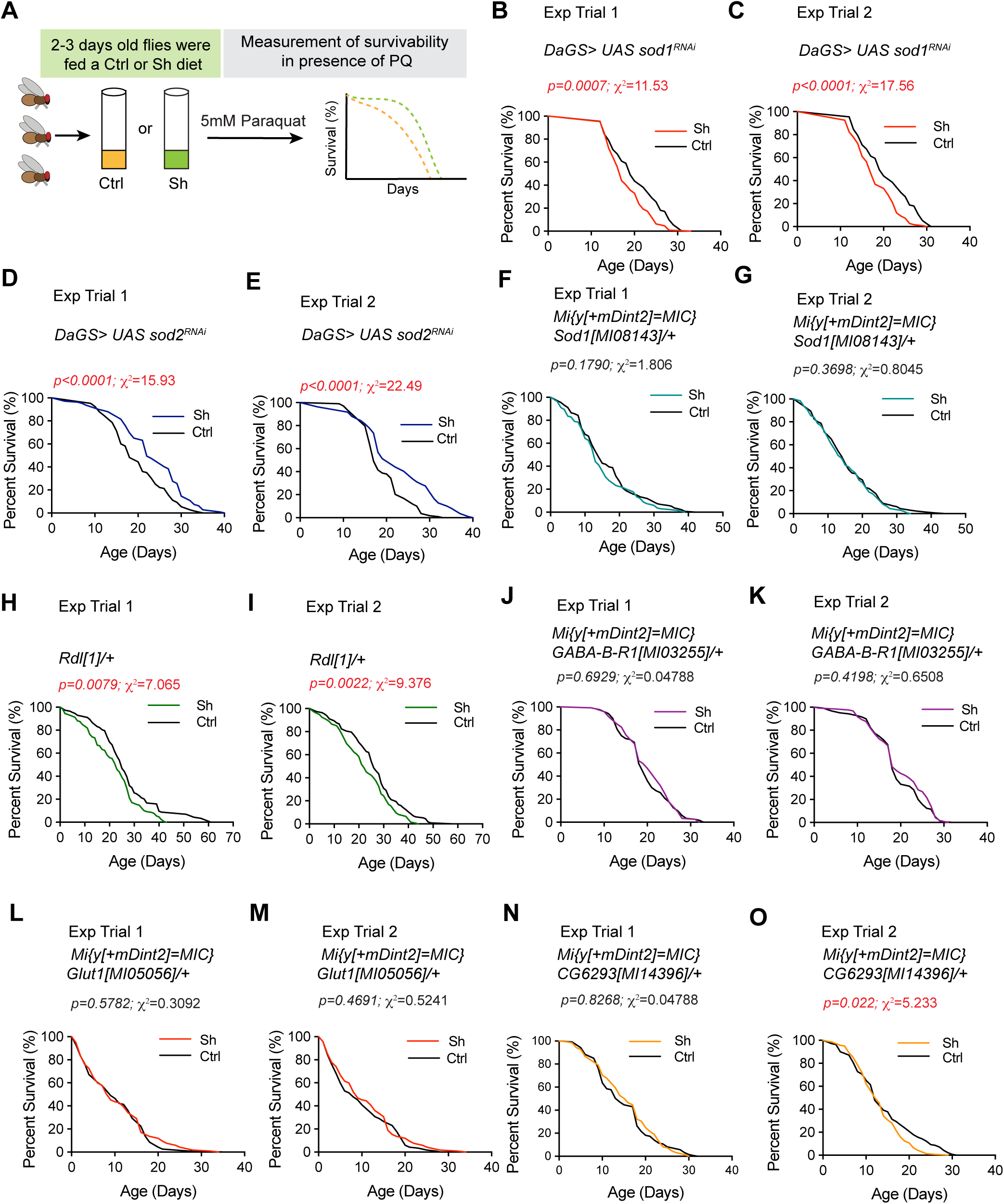
*Sod1, rdl, GABA-B-R1,* and *glut1* are required for the protective effects of *Convovulus pluricaulis* (Shankhpushpi)-supplemented diet under paraquat-induced oxidative stress conditions. (A) Dietary scheme for the analysis of different lines that were fed 5 mM paraquat (PQ) in either a control diet or a Shankhpushpi (Sh)- supplemented diet during adulthood. Survivability of flies that were fed a control or Shankhpushpi-supplemented diet was measured in presence of 5mM paraquat (PQ). (B-Two experiments denoting the survivability of female *daGS>UAS Sod1^RNAi^* flies that were fed a control or Sh diet with 5 mM paraquat in adult stages. Ubiquitous knockdown of *sod1* resulted in the loss of the increased survivability in the presence of Sh in the PQ diet. A moderate but significant decrease in lifespan was observed upon knockdown of *sod1* in Sh diet in both experimental trials. (D-E) The ubiquitous knockdown of *sod2* does not influence the ability of Sh to enhance survivability in paraquat-induced oxidative stress conditions. The *daGS>UAS Sod2^RNAi^* flies that were fed Sh survive significantly longer as compared to *daGS>UAS Sod2^RNAi^*flies that were fed a control diet. (F-G) Reducing one functional copy of *sod1* also worsens the increase in survivability seen upon intake of Sh. No significant difference was observed for flies that were fed a control or Sh diet. (H-I) Reducing the dosage of *rdl* abolishes the antioxidant effect of Sh. Flies expressing one copy of *rdl* have a significant reduction in survivability on the Sh-diet as compared to the control diet in the presence of PQ. (J-O) Reducing one copy of *GABA-B-R1* (J-K), *glut1* (L-M) or *CG6293* (N-O) also abolished the ability of Sh to enhance lifespan in presence of PQ. For statistical comparison of survival curves, p-values and χ ^2^ were calculated with log rank test and are noted in the Figure panels and Table 1. The maximum lifespan and medium lifespan and number of flies are noted in Table 1. Genotypes used in the study: (B-C) *DaGS>UAS Sod1^RNAi^*: w[*];*P { w[ +mW.hs] =Switch1}DaGS*;P{w[+mC]=UAS-Sod1.RNAi.H}4; P{w[+mC]=UAS-Sod1.RNAi.H}4 (D-E) *DaGS>UASSod2^RNAi^*: *+/+; P {y[+t7.7] v [+t1.8] =TRiP.GL01015} attP40*/*P { w[ +mW.hs] =Switch1}DaGS*; (F-G) *y*[1] *w[*]; Mi{y[+mDint2]=MIC}Sod1[MI08143]/+*; (H-I) *Rdl*[1]*/+*; (J-K) *y*[1] *w[*]; Mi{y[+mDint2]=MIC}GABA-B-R1[MI03255]/+*; (L-M) y[1] w[*]; Mi{y [+mDint2] =MIC}Glut1[MI05056]/+; (N-O) y[1]w[*];Mi{y[+mDint2]=MIC}CG6293 [MI14396] /+

**Table 1.**
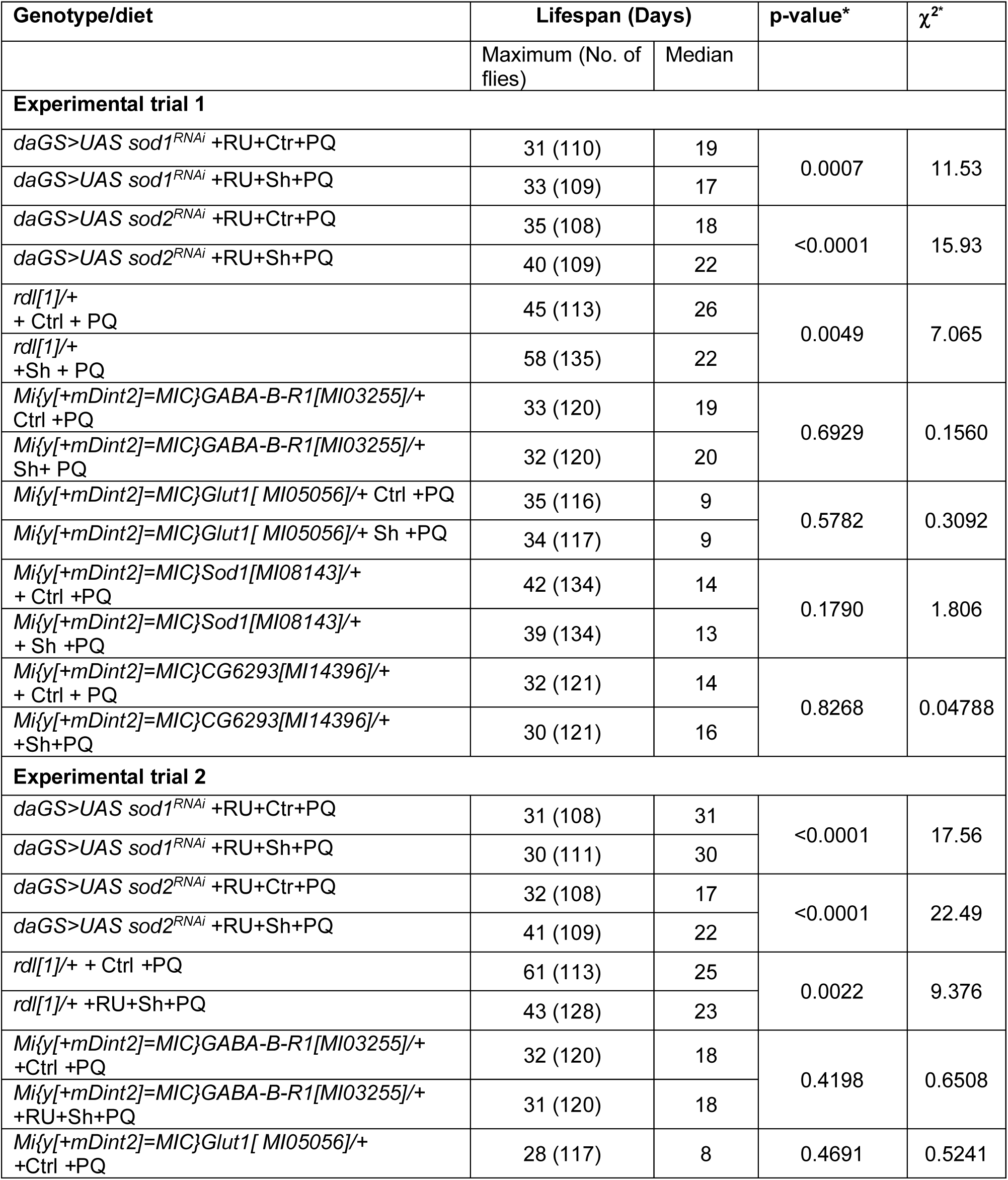
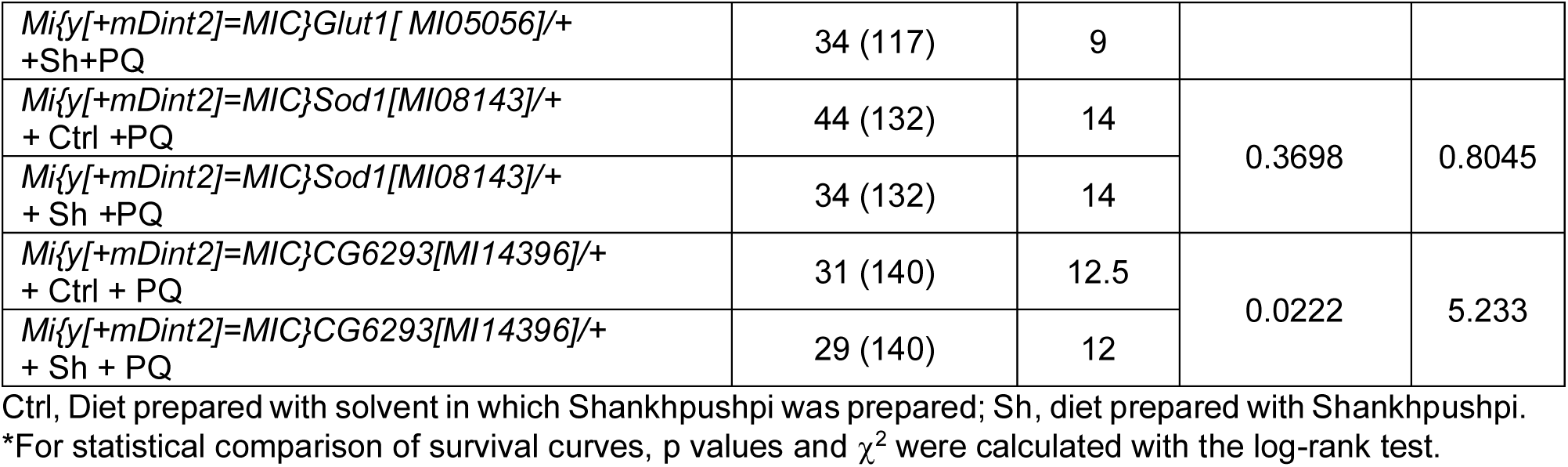
Survivability analysis of *rdl*, *sod1*, *GABA-B-R1*, *sod2*, *CG6293* and *glut 1* lines in paraquat diets.

## 4. DISCUSSION

### 4.1. Shankhpushpi regulates metabolic pathways associated with oxidative stress defense and brain function

*C. pluricaulis* (Shankhpushpi) plant extract has been extensively utilized in the pharmaceutical and nutraceutical industries as an antidepressant, anxiolytic and antioxidant [18, 22, 58, 60, 68–71]. Several studies have reported the phytochemical characterization of different plant parts, and other reports in animal models have confirmed its antioxidant effects [18, 24, 72–78]. Despite the extensive pharmacological characterization, the molecular players underlying the beneficial effects of this medicinal plant extract remain unknown. Here, we employed tissue-specific, untargeted metabolomic analysis of wild-type *Drosophila* fed a diet containing *C. pluricaulis* to investigate the metabolic changes induced by Shankhpushpi. Intake of *C. pluricaulis* (Shankhpushpi) induced significant changes in metabolites such as L-Ascorbate, Glucose, specifically in the head tissue and not in the body (**Figure 2 and Figures S2-3)**. This analysis was further validated by examining the expression of genes involved in the associated metabolic pathways (**Figure 3 and Figure 5**). Together, these data helped narrow down some key biological targets (*CG6293*, *sod1*, *rdl*, *GABA-B-R1*, and *glut1*).

The identification of these neuronally enriched effectors would not have been possible if the metabolomic analysis had been performed in whole animals instead of the tissue. The use of *Drosophila melanogaster* allowed us to narrow down the relevant targets by analyzing readily available mimic lines and RNAi lines. Thus, our approach highlights the utility and effectiveness of the genetically amenable fruit fly, *Drosophila melanogaster,* for illuminating the molecular mechanisms underlying natural dietary interventions.

### 4.2. Evidence for the role of L-Ascorbate and Ascorbate transporter in the management of depression-like behavior by *C. pluricaulis*

L-Ascorbate is the physiological anionic form of ascorbic acid or Vitamin C. Humans and insects are unable to synthesize ascorbate, and are dependent on dietary intake of Vitamin C. L-Ascorbic acid is a key cellular antioxidant in the central nervous system and plays a role in remodeling the epigenome by promoting ten-eleven translocation (TET) enzyme-mediated DNA oxidation[79–83]. Evidence from animal and human studies also indicates that psychosocial stressors lead to depression by driving epigenetic changes such as altered DNA methylation[84, 85]. Converging evidence has also revealed that oxidative stress is a key player in the pathogenesis of major depressive disorder[86, 87]. Oxidative stress leads to the oxidation of ascorbic acid, resulting in the generation of dehydroascorbic acid and an increased turnover of ascorbic acid. These findings from multiple studies suggest that disrupted ascorbic acid homeostasis may be responsible for the alterations in DNA methylation with oxidative stress under depression. Ascorbic acid homeostasis is regulated by two isoforms of the sodium-dependent vitamin C transporters (SVCT1 and SVCT2), which are coded by the two solute carrier family 23 members 1 and 2 genes, SLC23A1 and SLC23A2. SVCT 1 functions as the predominant carrier of ascorbic acid in the intestine and SVCT2 is primarily responsible for the transport of Ascorbic acid in the brain[43]. Previous studies have reported that oral administration of ascorbic acid can increase the fluoxetine antidepressant efficacy as an adjunct [88]. Pretreatment with a low dose of ascorbic acid has been shown to prevent chronic stress-induced depression like behavior[89, 90]. Several preclinical studies that have tested the use of Ascorbate in managing Major Depressive disorder (MDD) provide support for our work in *Drosophila* [41, 91]. These studies have utilized metabolomic approaches, treatment with inhibitors and polymorphisms in gene coding for ascorbate transporters such as SVCT1. Some of the mechanisms proposed for the anti-depressant action of L-Ascorbate include: (i) activation of serotonin, dopamine and beta-adrenergic receptors[91]; (ii) inhibition of nitric oxide production when NMDA receptors are activated[92, 93]; (iii) modulation of glutaminergic signaling[94]; and (iv) interference with transcription and activation of antioxidant enzymes[95]. According to Flybase (FB2025_2), the *Drosophila* gene *CG6293* has been predicted to be involved in the transmembrane transport of L-Ascorbic acid and is orthologous to SLC23A1(solute carrier family 23 member 1) and SLC23A2 (solute carrier family 23 member). Our metabolomic analysis in brain tissue indicates that L-Ascorbic acid levels in the brain tissue are enriched upon intake of Shankhpushpi, and the ubiquitous knockdown of *CG6293* abolished the anti-depressant effects of *C. pluricaulis* extract (Shankhpushpi) (Figure 4). By identifying a previously unknown link between *C. pluricaulis* intake and the expression and/or activity of Ascorbate transporter, we have uncovered one mechanism by which *C. pluricaulis* (Shankhpushpi) exerts its effect as an antidepressant. In addition, our study has also uncovered *sod1*, *GABA-B-R1*, *rdl*, and *CG6293* as molecular effectors of the antioxidant effect of *C. pluricaulis* (Shankhpushpi). Taken together, our findings have uncovered key conserved downstream molecular targets responsible for the neuropharmacological effects of *C. pluricaulis*. Our work highlights the genetically amenable fruit fly, *Drosophila melanogaster,* as an ideal model for evaluating the antidepressant efficacy of natural medicinal extracts. Future strategies aimed at developing and evaluating combinations of natural plant-based extracts will likely aid in designing an optimal supplement for the prevention and/or treatment of Major depressive disorder.

## CREDIT AUTHORSHIP CONTRIBUTION STATEMENT

**Shreyasi Mitra:** Writing-original draft, Writing-review & editing, Visualization, Validation, Methodology, Investigation, Formal analysis, Data curation. **Amit Kumar:** Writing-review & editing, Visualization, Validation, Methodology, Investigation, Formal analysis, Data curation. **Manish Pandey:** Visualization, Validation, Methodology, Investigation, Formal analysis, Data curation. **Meenakshi Sharma:** Investigation, Data acquisition. **Shivani Pundir:** Investigation, Data acquisition. **Mansi Jangir:** Investigation, Data acquisition.

**Geetanjali Chawla:** Writing-original draft, Writing-review & editing, Visualization, Validation, Supervision, Resources, Project administration, Methodology, Investigation, Funding acquisition, Formal analysis, Data curation, Conceptualization.

## ACKNOWLEDGEMENTS

We thank the Fly Facility at the Department of Life Sciences, Shiv Nadar Institution of Eminence, for their continuous support. Sakshi Bansal and Haseeb Ul Arfin for their support during the initial stages of this study. We are also grateful to the Regional Centre for Biotechnology, NCR Biotech Science cluster, and Shiv Nadar Institution of Eminence for providing the infrastructural support during the different stages of the study. Metabolomics analysis was performed at the Metabolomics Core Facility at the University of Utah. Mass spectrometry equipment at the University of Utah was obtained through NCRR Shared Instrumentation Grant 1S10OD016232-01, 1S10OD018210-01A1 and 1S10OD021505-01. We thank the Bloomington *Drosophila* Stock Center (NIH P40OD018537) and VDRC for providing fly stocks, as well as Flybase (NIH5U41HG000739). This research was supported by the DBT/Wellcome Trust India Alliance Fellowship/Grant [grant number IA/I(S)/17/1/503085] to G.C. and Shiv Nadar Institution of Eminence fellowship to S.M. and A.K.

## SUPPLEMENTARY FIGURES

**Figure S1:**
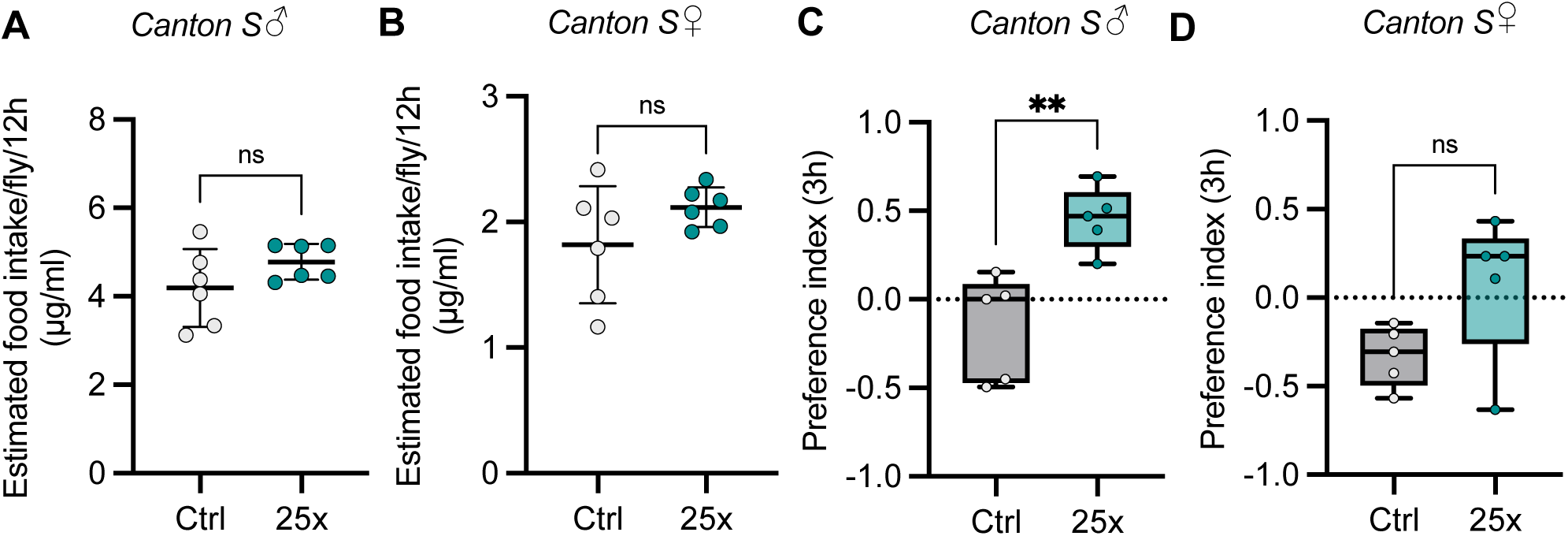
25X Shankhpushpi administration throughout development ameliorates stress-induced anhedonia in male *Canton S* flies but not females. (A-B) Male (A) and female (B) *Canton S* flies consume comparable amounts of normal fly food and 25X Shankhpushpi supplemented food. Food intake was measured by Ex-Q assay. Each data point represents a group of 10 flies. Data are represented as mean ± SD, n=6. P-value was calculated using an Unpaired t-test with Welch’s correction. (C-D) Canton S flies were raised on 25x Shankhpushpi supplemented fly food after which 3-5 days old flies were exposed to stress regime and their preference index was assayed using sucrose preference test. Male flies (C) showed positive sucrose preference index in comparison to regular food-fed stressed controls indicating relief from anhedonia due to Shankhpushpi administration while preference index of female flies (D) remained unchanged in comparison to their control flies. The box plot shows the median and the box spans from the first quartile to the third quartile. The whiskers extend from the box to the minimum and maximum data points, and we used an α level of 0.05 to assess statistical significance. *p<0.05, ***p<0.005 by Student t-test.

**Figure S2.**
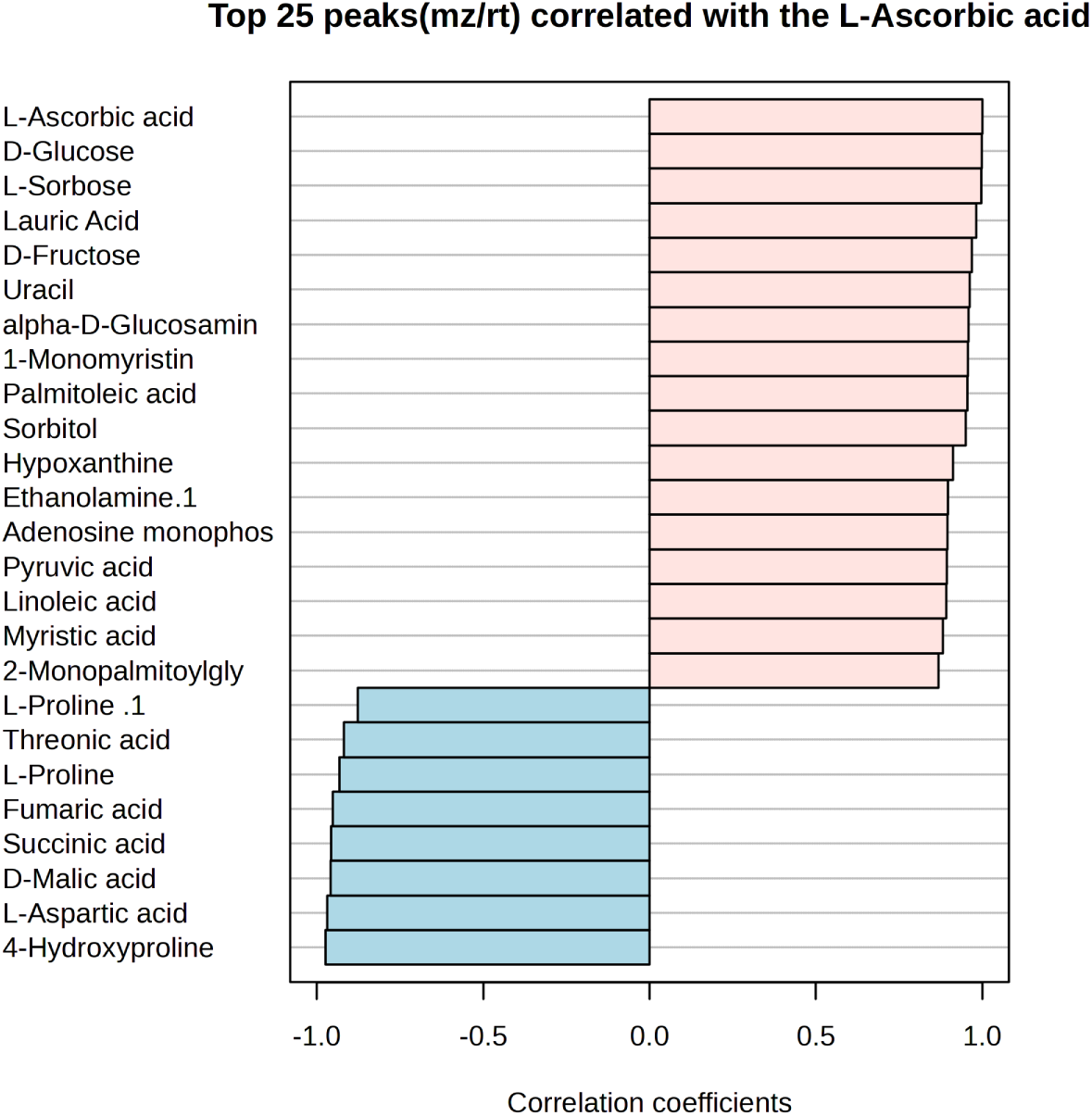
Correlation analysis of metabolites associated with L-Ascorbate. Top 25 metabolites correlated with L-Ascorbate levels across samples.

**Figure S3.**
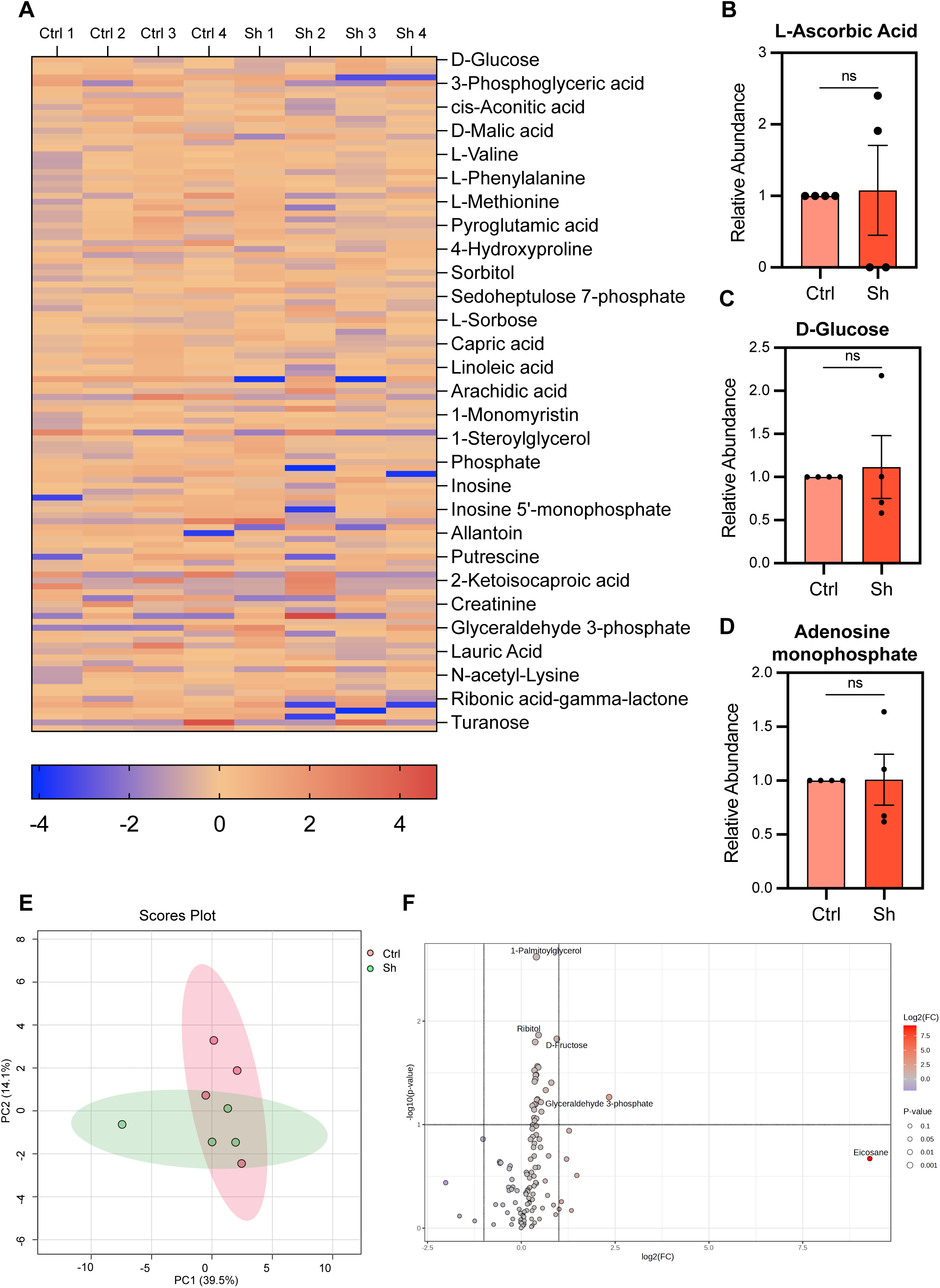
Metabolites in body tissue of Shankhpushpi-fed flies show no significant differences from controls. (A) Heatmap showing altered metabolites in the body tissue of Shankhpushpi-fed wild type flies in comparison to flies raised on control diet. Metabolomics analysis was carried out using four biological replicates for control and Shankhpushpi-fed group. (B-D) L-ascorbic acid (B), adenosine monophosphate(C), and D-glucose (D) levels remain unaffected in the body tissue of Shankhpushpi-fed flies in comparison to controls. (E) Projection of Shankhpushpi and control metabolite samples obtained from body tissue onto principal components 1 and 2 (PC1 vs PC2), explaining 39.5% and 14.1% of variance, respectively. Red points indicate regular fly food-fed controls and green points indicate metabolites in shankhpushpi-fed flies. (F) Volcano plot showing only D-fructose, glyceraldehyde-3-phosphate and Eicosane increased in body tissue of Shankhpushpi-fed flies compared to control flies, while no other metabolites differ significantly. Error bars indicate mean ± S.D. *p<0.05 by Student t-test.

**Figure S4.**
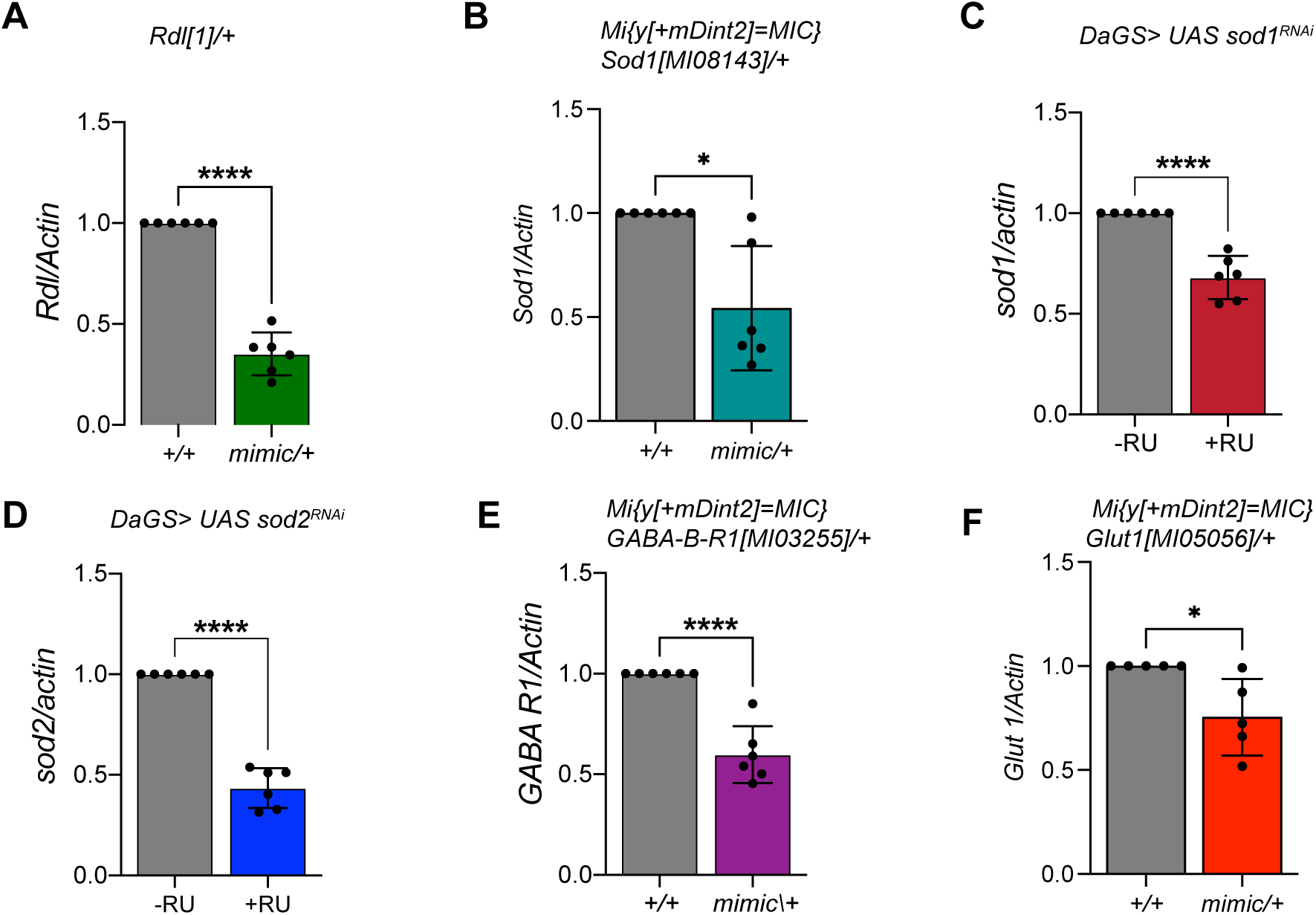
Validation of the knockdown of expression of *sod1*, *sod2*, *GABA-B-R1*, *rdl* and *glut1* in the lines that were used for survivability assays in presence of Paraquat-induced oxidative stress. (A) Quantitative RT-PCR of *rdl* with RNA extracted from 3–5-day-old w[1118] (+/+) and *rdl*[1]*/+* female flies. (B) Quantitative RT-PCR of *sod1* with RNA extracted from 3-5 day old *w*[1118] *(+/+)* and *Mi{y[+mDint2]=MIC} Sod1 [MI08143]/+* female flies. (C) A transgene expressing dsRNA for RNAi of *Sod1* was expressed ubiquitously using the steroid (RU-486) inducible gene switch Daughterless Gal4 driver. Quantitative RTPCR of *sod1* with RNA extracted from *daGS>UAS Sod1^RNAi^* female flies in presence or in absence of RU-486 for 5 days. (D) A transgene expressing dsRNA for RNAi of *Sod2* was expressed ubiquitously using the steroid (RU-486) inducible gene switch Daughterless Gal4 driver. Quantitative RTPCR of *sod2* with RNA extracted from *daGS>UAS Sod2^RNAi^* female flies in presence or in absence of RU-486 for 5 days. (E) Quantitative RT-PCR of *GABA-B-R1* with RNA extracted from 3–5-day-old w[1118] (+/+) and *Mi{y[+mDint2]=MIC}GABA-B-R1[MI03255]/+* female flies. (F) Quantitative RT-PCR of *glut1* with RNA extracted from 3-5 day old *w*[1118] *(+/+)* and Mi{y[+mDint2]=MIC} Glut1[MI05056]/+female flies.The expression of all the mRNAs tested was significantly reduced in the mimic lines (A-B and E-F) and upon RNA interference (C-D). Expression levels were normalized to *actin*. Data are represented as mean ± SD, n = 3. Two technical replicates were used for each biological replicate (n = 3) in the analysis. P-value was calculated using an unpaired t-test with Welch’s correction and is noted in the bar graph.

